# Quantitative mapping of autophagic cargo during nutrient stress reveals YIPF3-YIPF4 as membrane receptors for Golgiphagy

**DOI:** 10.1101/2022.12.06.519342

**Authors:** Kelsey L. Hickey, Sharan Swarup, Ian R. Smith, Julia C. Paoli, Joao A. Paulo, J. Wade Harper

## Abstract

During nutrient stress, macroautophagy is employed to degrade cellular macromolecules, thereby providing biosynthetic building blocks while simultaneously remodeling the proteome. While the machinery responsible for initiation of macroautophagy is well characterized, our understanding of the extent to which individual proteins, protein complexes and organelles are selected for autophagic degradation, and the underlying targeting mechanisms is limited. Here, we use orthogonal proteomic strategies to provide a global molecular inventory of autophagic cargo during nutrient stress in mammalian cell lines. Through prioritization of autophagic cargo, we identify a heterodimeric pair of membrane-embedded proteins, YIPF3 and YIPF4, as receptors for Golgiphagy. During nutrient stress, YIPF4 is mobilized into ATG8-positive vesicles that traffic to lysosomes as measured via Golgiphagy flux reporters in a process that requires the VPS34 and ULK1-FIP200 arms of the autophagy system. Cells lacking YIPF3 or YIPF4 are selectively defective in elimination of Golgi membrane proteins during nutrient stress. By merging absolute protein abundance with autophagic turnover, we create a global protein census describing how autophagic degradation maps onto protein abundance and subcellular localization. Our results, available via an interactive web tool, reveal that autophagic turnover prioritizes membrane-bound organelles (principally Golgi and ER) for proteome remodeling during nutrient stress.

**One-Sentence Summary:** During nutrient stress, macroautophagy uses organelle-phagy receptors to prioritize recycling of Golgi and ER membrane proteins.

## INTRODUCTION

Mammalian cells remodel their proteomes in response to changes in nutrient stress through transcriptional, translational, and degradative mechanisms (*1*). Central to these responses are proteasomal and autophagy-dependent degradative mechanisms that remove superfluous or damaged organelles and proteins to allow recycling of building blocks for cellular remodeling (*2*). Macroautophagy in response to reduced mTOR activity during nutrient stress has long been considered to result in non-specific capture of bulk cytoplasmic contents within autophagosomes, the biogenesis of which is dependent upon the VPS34 PI3P kinase, ULK1-FIP200 kinase complex, and core autophagy proteins (ATG9A, WIPI1/2, and the ATG8 lipidation machinery among others) (*3, 4*). However, recent work has revealed that selective forms of endoplasmic reticulum (ER) degradation by autophagy may be “hard-wired” into a broad autophagic response to nutrient stress (*5–10*). With ER-phagy, multiple partially redundant transmembrane ER proteins function as receptors to recruit core autophagy machinery, including the ULK1-FIP200 kinase complex (*6*), to initiate phagophore biogenesis proximal to the ER membrane (*10*). LC3-interaction regions (LIRs) within these receptors associate with the LIR docking site (LDS) in lipidated ATG8 proteins (6 orthologs in humans – MAP1LC3A, B, C and GABARAP, L1, L2) to facilitate ER engulfment within the phagophore (*10*).

Beyond ER-phagy, we have a limited understanding of the extent to which specific cargo are selected during macroautophagy and how cargo specificity is achieved. For example, widely studied ubiquitin-binding cargo adaptors that function in recognition of ubiquitylated autophagic cargo appear to play limited roles in cargo selection during nutrient stress, although two such receptors (NBR1 and TAX1BP1) have been linked with endosomal targeting and lysosomal degradation via the endosomal sorting complexes required for transport (ESCRT) system (*11, 12*). As such, several questions have emerged that are central to the field: 1) What proteins, protein complexes, and organelles are susceptible to autophagic degradation during nutrient stress? 2) Are there additional pathways for selective cargo degradation within the macroautophagy program and if so, how are these programs controlled? 3) How does the fraction of protein molecules degraded by autophagy scale with the total abundance of that protein within the cell and across individual sub-cellular compartments? In short, how selective is macroautophagy? Here, we combine quantitative global proteomics in autophagy proficient and deficient cells, ATG8-driven proximity biotinylation, and absolute protein abundance measurements to systematically map proteome and organelle responses to nutrient stress across sub-cellular compartments. Using these approaches coupled with an autophagic cargo prioritization scheme, we have generated a high confidence autophagy factor database, and through this, identified a pathway for selective autophagy of Golgi membrane proteins. Additionally, we have developed a ‘protein census’ describing how autophagic degradation maps onto protein abundance and sub-cellular localization across the proteome, to comprehensively define the selectivity of macroautophagy during nutrient stress.

## RESULTS

### Orthogonal proteomic approaches for Organelle-phagy receptor identification

Our previous studies using global proteome abundance analysis in HEK293, HEK293T, and HCT116 cells during amino acid (AA) withdrawal or MTORC1 inhibition with the small molecule inhibitor Torin1 (Tor1) revealed autophagy-dependent loss of on average ~8% of ~310 ER proteins (**Figure 1A**), but with little evidence of autophagy-dependent reduction in proteins from other compartments. (**fig. S1A**) (*5, 13*). Interestingly, AA withdrawal or MTORC1 inhibition resulted in ~10% reduction in Golgi membrane and associated proteins as annotated in Uniprot and this was blocked in cells lacking ATG7 or FIP200 (also called RB1CC1), components of the autophagy conjugation and initiation machinery, respectively (**Figure 1A and fig. S1A**). While multiple membrane-embedded autophagy receptors that are selective for turnover of ER by autophagy have been reported, membrane-embedded proteins that are selective for Golgi turnover by autophagy are lacking (*5–10, 14, 15*).

**Figure 1.**
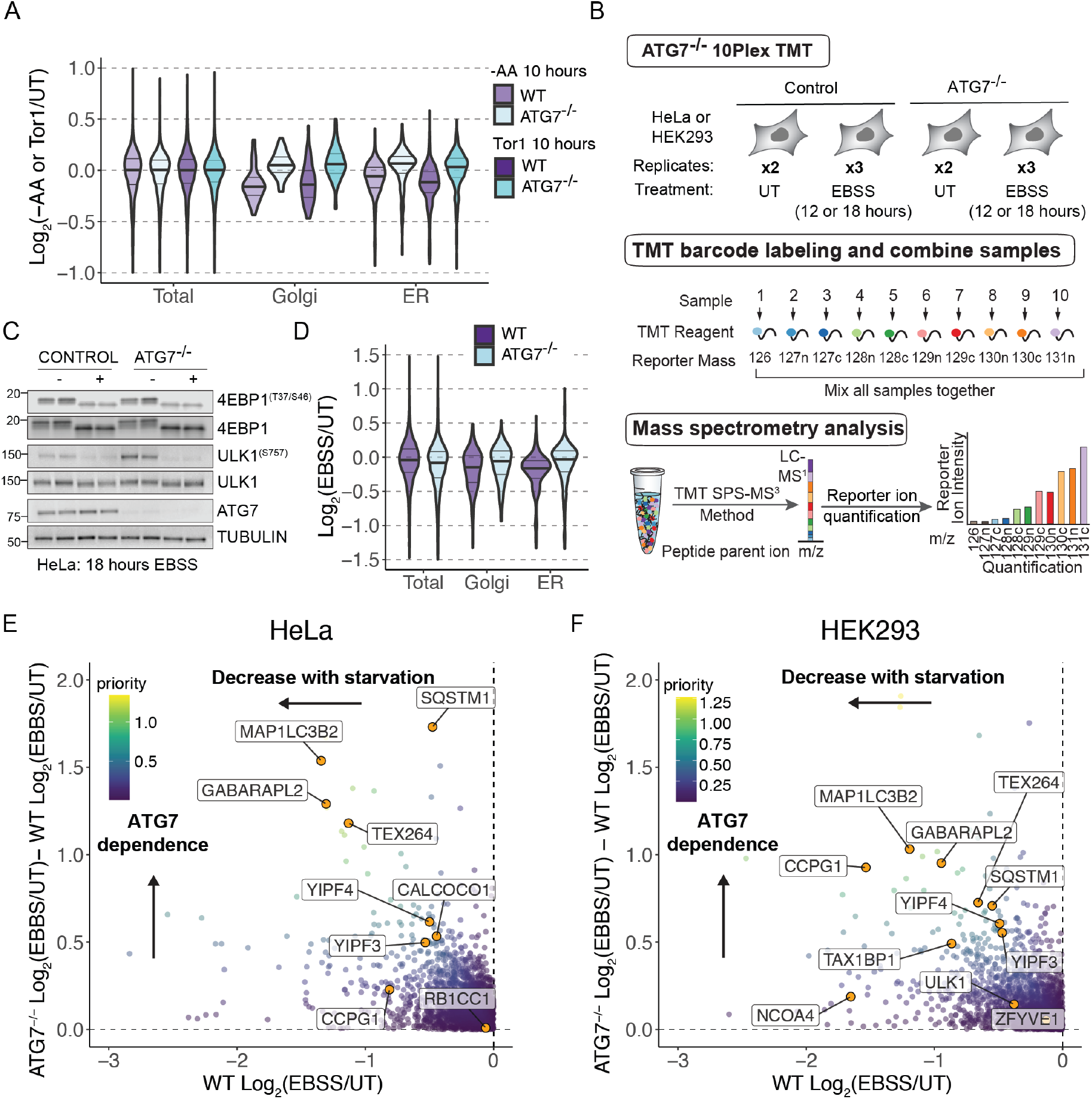
Global proteomic analysis of autophagy-dependent proteome remodeling with nutrient stress. (**A**), Violin plots of relative total (8,032, 8,412), Golgi (56, 60), and ER (340, 349) protein abundance in response to amino acid withdrawal or Torin1 treatment (10 hours) in WT or ATG7^-/-^ HEK293 cells from our previous study. (**B**) The 10-plex TMT pipeline to measure relative protein abundance during nutrient stress with or without active autophagy. Normalized total cell extracts were processed for 10-plex TMT mass spectrometry (TMT-MS) experiments. EBSS, withdrawal of amino acids and serum; UT, untreated. (**C**) Western blot showing markers of starvation (ULK1, 4EBP dephosphorylation) and ATG7 in WT and ATG7^-/-^ HeLa cells grown in EBSS for 18 hours. (**D**) Violin plots of relative total (8258), Golgi (160), or ER (344) protein abundance in response to EBSS treatment (12 hours) in WT and ATG7^-/-^ HeLa cells. (**E, F**) Plots of ATG7^-/-^ Log_2_(EBSS/UT) – WT Log_2_(EBSS/UT) versus WT Log_2_FC (EBSS/UT) for HeLa cells (panel E) and ATG7^-/-^ Log_2_(EBSS/UT) – WT Log_2_(EBSS/UT) versus WT Log_2_FC (EBSS/UT) for HEK293 cells (panel D) where priority for individual proteins is scaled based on the color code inset. Full plots are shown in **fig. S1G, H**.

To search for candidate Golgiphagy receptors during nutrient stress, we employed two approaches in parallel: 1) global quantitative proteomics with and without ATG7 in the same tandem mass tagging (TMT) plex in order to directly reveal autophagy-dependent changes in protein abundance, and 2) ATG8- driven proximity biotinylation (*16*). Our expectation, based in part on the behavior of ER-phagy receptors (*5*), is that candidate Golgiphagy receptors would decrease in abundance to a greater extent than Golgi proteins broadly, allowing their identification by global proteomics. In addition, we expect that relevant receptors would be in proximity with ATG8 proteins during the autophagy process, in an LDS dependent manner, allowing their identification by time-resolved proximity biotinylation with APEX2-ATG8 proteins (*5*).

HEK293 or HeLa cells with or without ATG7 were subjected to 12 or 18 hours of nutrient stress (EBSS), respectively, followed by analysis by 10-plex TMT proteomics (**Figure 1B, C, fig. S1B, Data S1**). Consistent with previous studies, proteins localized in ER and Golgi were reduced in an autophagy-dependent manner (**Figure 1D**). To identify candidate autophagy receptors and substrates, we calculated the starvation and ATG7 dependent decrease in protein levels (**Figure 1E, F**). As expected, we observed reduced levels of several selective autophagy receptors (TEX264, CCPG1, and SQSTM1) as well as ATG8 proteins MAP1LC3B and GABARAPL2 in one or both cell lines and this reduction was dependent on ATG7 (**Figure 1E, F, fig. S1C-H**) (*5, 13*). Interestingly, two Golgi proteins – YIPF3 and YIPF4 – stood out as proteins whose abundance was substantially reduced during starvation and the extent of dependence on ATG7 was comparable to other well-validated receptors (**Figure 1E, F, fig. S1G, H**).

We next employed proximity biotinylation in ATG8 knockout HeLa cells (*17*) that were reconstituted with APEX2-ATG8 proteins (MAP1LC3B^-/-^ cells reconstituted with APEX2-MAP1LC3B or GABARAPL2^-/-^ cells reconstituted with APEX2-GABARAPL2) (**Figure 2A, B**). To facilitate the identification of autophagy receptors associate with ATG8 proteins, we also performed analogous experiments with LDS mutations in GABARAPL2 (Y49A/L50A) or LDS and lipidation mutations in MAP1LC3B (K51A/G120A) (*18*). Cells were left untreated or subjected to nutrient stress (EBSS+BafilomycinA, (BafA)) for 3 hours prior to proximity biotinylation and proteomic analysis using 10-plex TMT (**Figure 2B, Data S2**). BafA treatment blocks lysosomal acidification, thereby blocking degradation of biotinylated proteins captured by autophagy and delivered to the lysosome. Among the most enriched proteins for both ATG8s in response to nutrient stress were ubiquitin-binding autophagy receptors (TAX1BP1, CALCOCO2, CALCOCO1, SQSTM1), ER-phagy receptors (CCPG1, TEX264), and core autophagy machinery (WIPI2, ULK1, FIP200/RB1CC1, and ATG8 proteins) (**Figure 2C, D, fig. S2A, B**). This enrichment was dependent on the LDS for known autophagy receptors (**Figure 2C, D, fig. S2C-F**).

**Figure 2.**
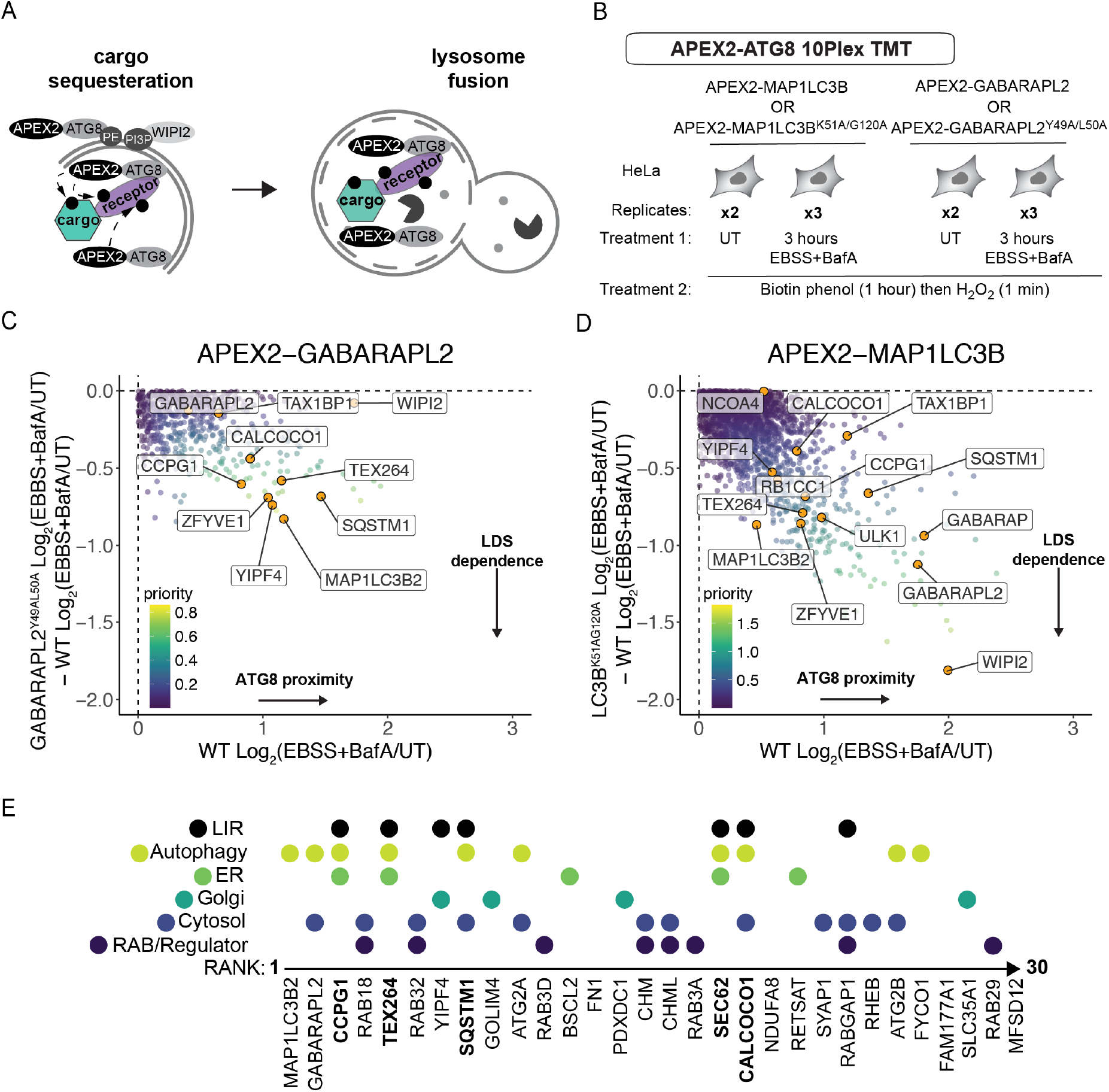
ATG8-drive proximity biotinylation identifies candidate Golgiphagy receptors. (**A**) Scheme depicting unbiased identification of ATG8-interacting proteins by proximity biotinylation. APEX2-ATG8 proteins are concentrated on the phagophore during the response to nutrient stress and function to recruit autophagic cargo receptors, which can be biotinylated by APEX2 during cargo engulfment. (**B**) 10-plex TMT APEX2-ATG8 pipeline to capture autophagy receptors during nutrient stress (EBSS + BafA, 4 hours) with or without active LIR docking sites. At 3 hours post-nutrient stress, cells were supplemented with biotin phenol (1 hour) and then treated with H_2_O_2_ for 1 min followed by quenching (see METHODS). (**C, D**) Plots of GABARAPL2^Y49A/L50A^ Log_2_(EBSS+BafA/UT) – WT Log_2_(EBSS+BafA/UT) versus WT Log_2_(EBSS+BafA/UT) (panel C) and MAP1LC3B^K51A/G120A^ Log_2_(EBSS+BafA/UT) – WT Log_2_(EBSS+BafA/UT) versus WT Log_2_(EBSS+BafA/UT) (panel D) where priority for individual proteins is scaled based on the color code inset. Full plots are shown in **fig. S2E, F**. (**E**) Top ranked proteins (n=30) based on summed individual rankings for global proteomics and ATG8 proximity biotinylation (see METHODS) displayed based on their sub-cellular localization, involvement in autophagy, and the presence of a known or candidate LIR motif. Proteins known to function as autophagic cargo receptors are in bold font.

To generate a prioritized collection of candidate autophagic factors, we first independently ranked proteins based on the extent of autophagic and starvation-dependent turnover for total proteome. Then we ranked proteins based on their LDS dependent ATG8 interaction from our APEX2 experiments (see METHODS). We then summed the individual rankings to generate a composite ranking and further classified proteins based on the presence of a previously identified or predicted LIR motif (**Figure 2E**, **Data S3**). The utility of this approach is indicated by the presence of TEX264, CCPG1, SQSTM1 and two ATG8 proteins within the top 10 ranked proteins (**Figure 2E**). The highest ranked Golgi protein (ranked 7^th^) was YIPF4 (**Figure 2E**). It was also the most strongly enriched Golgi protein with GABARAPL2 and its proximity biotinylation was dependent on a functional LDS (**Figure 2C, fig. S2A, C, E**). With APEX2- MAP1LC3B, YIPF4 displayed less enrichment but was nevertheless partially dependent on a functional LDS (**Figure 2D, S2B, D, F**). In addition, YIPF4 has a predicted LIR motif (**Figure 2E**) (*19*), making it a top candidate to be a Golgiphagy receptor.

### LIR-containing YIPF3/4 are in proximity with autophagy machinery during nutrient stress

YIPF3 and 4 are members of a family of Golgi proteins that each contain 5 transmembrane segments, with a cytosolic N-terminal region and a lumenal C-terminal region (**Figure 3A**). Although poorly studied, a previous report indicates that YIPF3 and YIPF4 can co-immunoprecipitate and are thought to form heterodimers (*20*). ColabFold implementation of AlphaFold (*21*) predicts the formation of a heterodimer involving interaction between transmembrane helices 1 (G155-T173) and 2 (M184-L208) in YIPF3 and transmembrane helices 2 (R135-V157) and 5 (L225-T242) in YIPF4, with both N-terminal regions being largely unstructured (**Figure 3B**). Deletion of YIPF4 in HeLa cells resulted in loss of YIPF3 (**Figure 3C**), indicating that YIPF3 stability likely requires association with YIPF4. Importantly, both YIPF3 and YIPF4 contain candidate LIR motifs in their cytosolic N-terminal tails, making them accessible to interactions with ATG8 proteins (**Figure 3A, B**).

**Figure 3.**
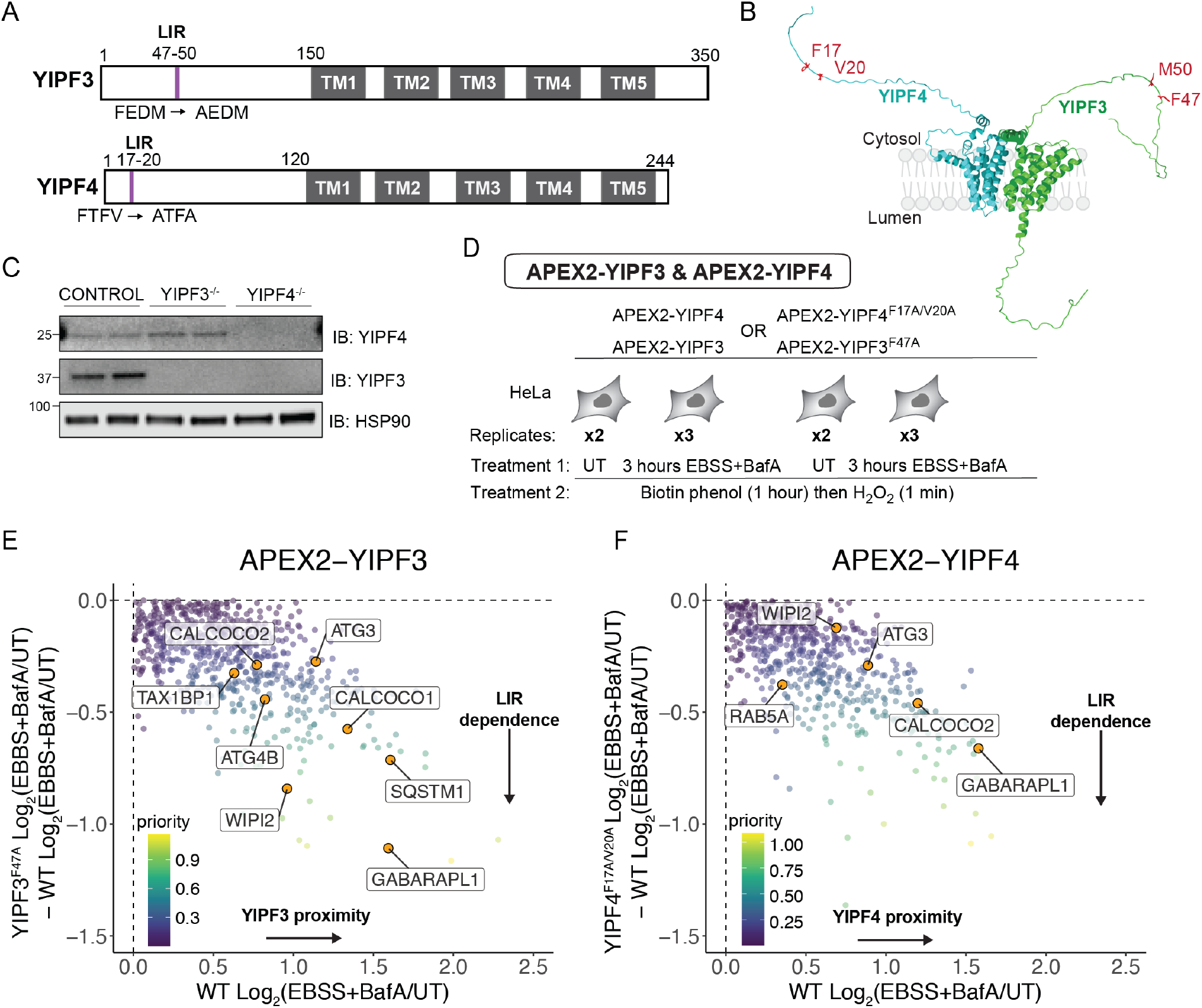
LIR-containing YIPF3 and YIPF4 associate with autophagy machinery during nutrient stress. (**A**) Domain structure of YIPF3 and YIPF4 showing the location of the five transmembrane segments and N-terminal candidate LIR motifs. (**B**) Model for YIPF3-YIPF4 heterodimer was generated using Colabfold. (**C**) Immunoblotting of WT, YIPF3^-/-^, and YIPF4^-/-^ HeLa cells probed in duplicate with the indicated antibodies. Anti-HSP90 was used as a loading control. (**D**) Experimental scheme for proximity biotinylation using APEX2-YIPF3/4 (or LIR mutants) in response to nutrient stress (EBSS, 4 hours). (**E**) Plots of YIPF3^F47A^ Log_2_(EBSS+BafA/UT) – WT Log_2_(EBSS+BafA/UT) versus WT Log_2_(EBSS+BafA/UT) (left) and YIPF4^F17A/V20A^ Log_2_(EBSS+BafA/UT) – WT Log_2_(EBSS+BafA/UT) versus WT Log_2_(EBSS+BafA/UT) (right) where priority for individual proteins is scaled based on the color code inset. Full plots are shown in **fig. S3E, F**.

To explore YIPF3/4 interactions during nutrient stress, we stably expressed APEX2-YIPF3 or APEX2-YIPF4 (**Figure 3A**) in HeLa cells lacking YIPF3 or YIPF4, respectively, and performed proximity biotinylation after 3 hours of nutrient stress (EBSS+BafA) (**Figure 3D, Data S4**). To determine LIR dependent interactors, we included APEX2 fusion proteins harboring mutations in the candidate LIR motifs for both YIPF3 and YIPF4 (**Figure 3A**). Among the most enriched proteins with wild type (WT) YIPF3 and YIPF4 was the ATG8 protein GABARAPL1 (**Figure 3E, F, fig. S3A, B**). These interactions were partially dependent on a functional LIR motif, indicating that YIPF3 and YIPF4 are in proximity to ATG8 proteins during nutrient stress and providing reciprocal validation of ATG8 proximity biotinylation described above (**Figure 3E, F, fig. S3C-F**). Additional autophagy factors, including WIPI1/2, ATG3 and ATG4B proteins, were also enriched, consistent with proximity to proteins involved in phagophore formation (**Figure 3E, F, fig. S3E, F)** (*3*).

### YIPF4-containing vesicles release from Golgi during nutrient stress via autophagy

Previous studies suggest that ER-phagy receptors promote ER capture via templating of phagophore formation on the ER membrane, with phagophore closure coupled to scission of the ER membrane (*22, 23*). To examine YIPF4 mobilization into vesicles during nutrient stress, we first created WT or FIP200^-/-^ HEK293 cells in which the endogenous N-terminus of YIPF4 was edited to append a monomeric neon green fluorescent protein (mNEON) (**fig. S4A**, see **METHODS**). mNEON-YIPF4 was localized to Golgi in untreated cells as indicated by extensive colocalization with the Golgi marker GOLGB1 (**Figure 4A, fig. S4B, C**). Strikingly, within 3 hours of starvation (EBSS+BafA), numerous mNEON-YIPF4-positive puncta (~500 nm in diameter) were observed (**Figure 4B, C, fig. S4B, C**). These puncta depended on the presence of BafA to block mNEON quenching and degradation within the lysosome (**Figure 4B, C**), and a subset of mNEON-YIPF4 puncta were found to co-localize with LAMP1, indicating trafficking to the lysosome (**Figure 4C, fig. S4B, C**). Importantly, the formation of these puncta was abolished in cells lacking FIP200 and in cells treated with a small molecule inhibitor of VPS34 (SAR405, VPS34i) (**Figure 4D, E, fig. S4B, C**), suggesting an essential role for autophagy in the liberation of YIPF4- positive vesicles from Golgi during nutrient stress, as is also seen with ER-phagy receptors (*22, 23*). Consistent with this, a subset of mNEON-YIPF4 puncta also co-localized with MAP1LC3B puncta, as assessed using immunofluorescence (**Figure 4F, fig. S4D**). As shown in **Figure 3E-F**, the ubiquitin-binding autophagy receptors SQSTM1, CALCOCO1, and CALCOCO2 (also called NDP52) were identified in APEX2-YIPF3/4 proximity biotinylation experiments. However, we found that addition of the ubiquitin E1 activating enzyme inhibitor TAK243 (*24*) had no effect on the liberation of mNEON-YIPF4 puncta in response to nutrient stress (**Figure 4G, fig. S4C**), suggesting the absence of a requirement for ubiquitin conjugation in this process.

**Figure 4.**
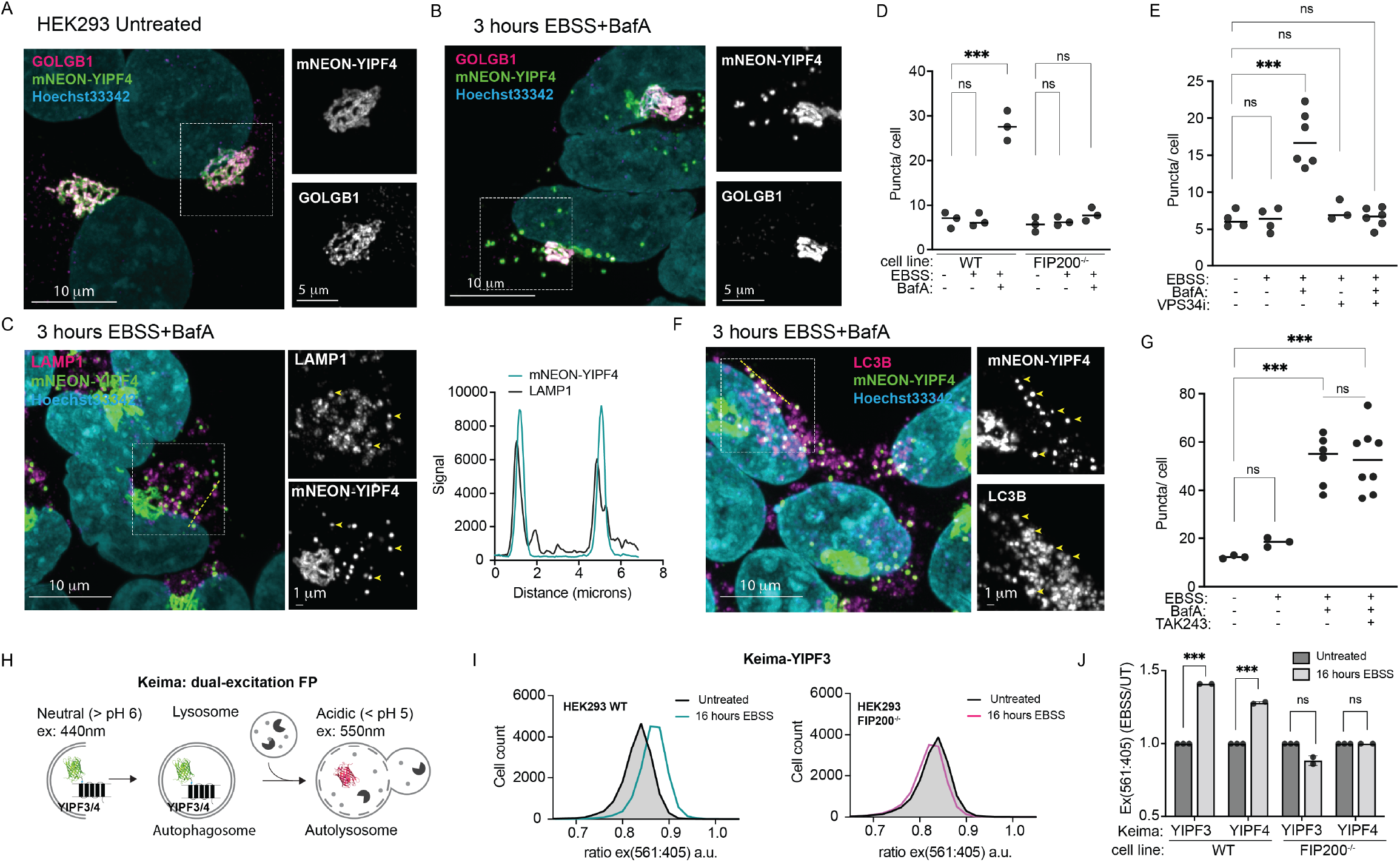
Golgi-derived YIPF4-containing vesicles traffic to lysosomes via autophagy. (**A, B**) HEK293 cells expressing endogenous YIPF4 tagged on its N-terminus with mNEON (green) by gene editing were imaged using confocal microscopy and co-stained with the Golgi marker GOLGB1 (magenta). Cells were either left untreated (panel **A**) or subjected to nutrient stress (3 hours) (panel **B**) prior to imaging. Nuclei were labeled with Hoechest33342 dye (cyan). Scale bars 5 or 10 microns as indicated. (**C**) Cells treated as in panel **B** were immunostained with a-LAMP1 (magenta). Line scans (right histogram) demonstrate colocalization of mNEON-YIPF4 puncta and LAMP1-positive structures. Scale bars 1 or 10 microns as indicated. Yellow arrowheads indicate examples of YIPF4-positive puncta that overlap LAMP1-positive structures. (**D, E**) The number of mNEON-YIPF4 puncta/cell is plotted for the indicated treatments in cells with or without FIP200. Each dot represents one image where the mNEON and nuclei were counted. ***, Mann-Whitney p-value < 0.05. (**F**) Cells treated as in panel **B** were immunostained with a-LC3B (magenta). Yellow arrowheads indicate examples of YIPF4-positive puncta that overlap LC3B-positive structures. Scale bars 1 or 10 microns as indicated. (**G**) The number of mNEON-YIPF4 puncta/cell is plotted for the indicated treatments in cells 3 hours post nutrient stress. ***, Mann-Whitney p-value < 0.05. (**H**) Scheme outlining Keima-YIPF3/4 as reporters for Golgiphagic flux. (**I**) Keima-YIPF3 expressing HEK293 cells (with or without FIP200) were left untreated or subjected to nutrient stress for 16 hours and then analyzed by flow cytometry. Frequency distributions of 561/405 nm ex. ratios are shown (*n* = 10,000 cells per condition). (**J**) Bargraph of median values of the biological duplicate experiments for 561/405 nm ex. ratios for Keima-YIPF3 or Keima-YIPF4 with or without FIP200. Error bars represent standard error of the mean (s.e.m.).

### Keima-YIPF3 and YIPF4 undergo autophagic flux and function as reporters of Golgiphagy

To examine Golgiphagic flux in HEK293 cells, we fused the fluorescent Keima protein to YIPF3 and YIPF4 (**Figure 4H**). Keima is a pH-responsive reporter that undergoes a change in chromophore resting state upon trafficking to the lysosome (pH of ~4.5), allowing flux measurements in single cells by determining the ratio of 561 nm/405 nm excitation via flow cytometry (*25*). We found that both Keima- YIPF3 and Keima-YIPF4 flux was increased upon nutrient stress and this flux was completely blocked in FIP200^-/-^ cells (**Figure 4I, J**). Thus, like other membrane-bound autophagy receptors (*5, 7*), Keima-YIPF3 and YIPF4 are Golgiphagy reporters that can report on Golgi trafficking to the lysosome in a manner that depends on core autophagy machinery.

### Quantitative mapping of autophagy-dependent degradation programs during nutrient stress

The finding that YIPF4-positive vesicles traffic through the autophagy system led us to explore in detail the identity of Golgi proteins that are degraded by autophagy during nutrient stress, and how this is integrated into global protein turnover via autophagy. We reasoned that autophagy receptors and cargo would all behave in a similar manner upon nutrient stress, i.e degraded when autophagy is intact, but not in autophagy deficient cells. To gain a comprehensive and unbiased understanding of all proteins that behave in a similar manner to well characterized autophagy clients during nutrient stress, we used a set of known autophagy proteins (e.g. ER-phagy receptors, ATG8 proteins, CALCOCO1), and calculated the median value (see **METHODS**) in each condition to construct a consensus profile using our data from **Figure 1** wherein WT or ATG7^-/-^ HEK293 cells were treated with or without EBSS for 12 hours (**Figure 5A, fig. S5A, B**). Next, to identify proteins displaying similar condition profile, we calculated the root-mean-square error (RMSE) for every protein quantified, across all treatments and replicates. This analysis is analogous to ‘protein correlation profiling’ (*26*). Proteins with lower total RSME more closely resemble the normalized abundance profile of known autophagy proteins and ideally should be enriched in receptors or clients of autophagy (**Figure S5A**). With this approach, we prioritized 732 proteins whose abundance profile is concordant with starvation and autophagy dependent turnover: decreased abundance with EBSS treatment that is blocked by deletion of ATG7 (**Figure 5B**). Golgi and ER proteins display lower RMSE compared to all other organelles, while proteins from cellular compartments such as the nucleus or ribosomal proteins displayed a high RMSE consistent with their non-involvement in starvation-based macroautophagy (**Fig. S5C, D**) (*13*). In parallel, we performed an orthogonal experiment using an autophagy mutant in a distinct branch of the pathway (FIP200^-/-^) in the context of an alternative nutrient stress (AA withdrawal) (**Figure 5A, fig. S5E, F**). For AA withdrawal, we prioritized 684 proteins, which had the lowest RMSE and displayed a profile consistent with autophagy and starvation-dependent turnover (**Figure 5C, fig. S5G-I**). We refer to these groups of proteins as candidate ‘autophagy’ proteins. We anticipate that these ‘autophagy’ candidates should prioritize proteins that are degraded by autophagy during nutrient starvation, potentially decoupling autophagy from other starvation dependent cellular responses (e.g. alternative pathways such as proteasome or ESCRT-dependent endolysosomal degradation or translation inhibition).

**Figure 5.**
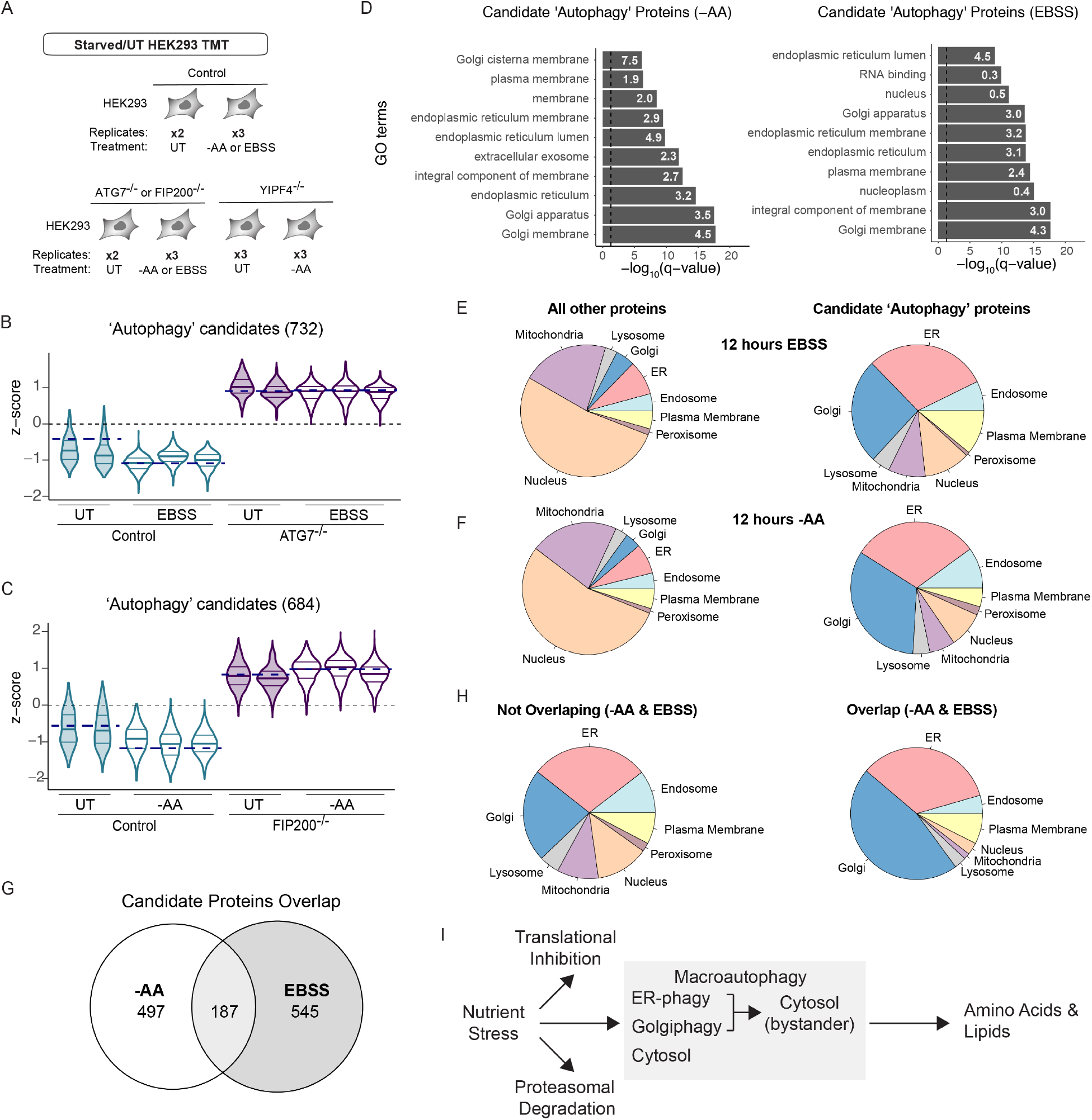
Golgi and ER represent major targets for autophagic turnover in response to nutrient stress. (**A**) Scheme depicting method for global proteome alterations via autophagy in response to nutrient stress. (**B**) Violin plots for proteins identified as candidate ‘autophagy’ proteins in WT and ATG7^-/-^ HEK293 cells with or without EBSS (12 hours). Navy dashed lines indicated median value for known autophagy proteins in each condition. (**C**) Violin plots for proteins identified as candidate ‘autophagy’ proteins in WT and FIP200^-/-^ HEK293 cells with or without AA withdrawal (12 hours). Navy dashed lines indicated median value for known autophagy proteins in each condition. (**D**) Top 10 Gene Ontology terms identified for candidate ‘autophagy’ proteins from cells subjected to EBSS treatment (left panel) or AA withdrawal (right panel). (**E**) Frequency of proteins with the indicated sub-cellular localizations for the candidate ‘autophagy’ proteins or all other proteins for cells subjected to EBSS treatment. (**F**) Frequency of proteins with the indicated sub-cellular localizations for the candidate ‘autophagy’ proteins or all other proteins for cells subjected to AA withdrawal. (**G**) Overlap of proteins identified in the candidate ‘autophagy’ proteins from both types of nutrient stress. (**H**) Frequency of proteins with the indicated sub-cellular localizations for either overlapping or non-overlapping proteins from panel G. (**I**) Model for how nutrient stress activates a macroautophagy program wherein ER and Golgi turnover by selective autophagy underlies a major component of the process. Cytosolic proteins are also degraded but could also be captured non-specifically as part of the selective autophagy program. See text for details.

Gene Ontology (GO) analysis of candidate ‘autophagy’ proteins from both types of nutrient stresses revealed dramatic enrichment in terms linked with ER and Golgi, which were prominently present in the top 10 terms identified (**Figure 5D**). We next compared these ‘autophagy’ candidate proteins to all other proteins quantified in the experiments and found that Golgi and ER were the most over-represented sub- cellular compartment across all compartments examined (**Figure 5E, F**). Upon closer examination of proteins within the Golgi, we found that this signature is primarily composed of membrane-embedded Golgi proteins (**fig. S6A**). Across the two independent experiments with distinct types of nutrient stresses, we identified 187 proteins in common in both sets of ‘autophagy’ candidate proteins (**Figure 5G**). The common proteins, compared with non-overlapping proteins, are even more over-represented in terms of sub-cellular localization within Golgi and ER (**Figure 5H**). Additionally, Golgi and ER display strong overlap of proteins compared with other compartments, including the cytosol, that, in turn, have a decreased proportional overlap between the two distinct nutrient stressors (**fig S6B, C**). By examining the enrichments of proteins with an autophagic turnover signature and their overlap in two independent experiments, our analysis revealed selective programs directed toward Golgi and ER within nutrient stress-dependent macroautophagy (**Figure 5I, see Discussion**).

### Selectivity of YIPF3/4-dependent Golgiphagy for Golgi-membrane proteins

The results described thus far suggest a role for YIPF4 in autophagic turnover of Golgi. To directly examine this possibility and to determine specific cargo, we included YIPF4^-/-^ HEK293 cells in the same TMT proteomics experiment examining FIP200-dependent cargo upon AA withdrawal (**Figure 5A, fig. S5D, Data S5**). YIPF4^-/-^ cells display loss of YIPF3, and therefore may mimic double knock-out cells, but still respond to AA withdrawal signaling as demonstrated by immunoblotting of cell extracts with anti-p- 4EBP1 or anti-p-ULK1 (**fig. S5D**). Since YIPF4 is a Golgi protein and Golgi is over-represented in the candidate ‘autophagy’ proteins, we initially examined the behavior of Golgi proteins among this ‘autophagy’ candidate cohort in cells lacking YIPF4. Golgi proteins fall into two major classes – those proteins that contain one or more integral transmembrane segments and those that are soluble proteins but spend part of their life-history in association with the Golgi – which we refer to as Golgi-membrane and Golgi-associated, respectively. We found that while loss of YIPF4 had no effect on degradation of non-Golgi proteins during AA withdrawal, YIPF4^-/-^ cells displayed an intermediate effect on the abundance of 79 Golgi proteins within the ‘autophagy’ candidate proteins (**Figure 6A**). The contribution of YIPF4 to Golgi turnover was found to be largely specific to Golgi-membrane proteins, with little effect on bulk Golgi-associated proteins (**Figure 6A**). The selectivity for Golgi-membrane proteins and the absence of a strong effect of YIPF4 deletion on the abundance of other sub-cellular compartments is indicated by correlation plots of YIPF4^-/-^ and FIP200^-/-^ cells with or without AA withdrawal (**Figure 6B, fig. S7A**). In particular, bulk ER protein abundance was stabilized in FIP200^-/-^ cells but unaffected in YIPF4^-/-^ cells, highlighting the specificity of YIPF4 for Golgiphagy (**Figure 6B**). Golgi ‘autophagy’ candidate proteins are displayed in **Figure 6C** based on their trans-membrane segment disposition, and the extent of stabilization by FIP200 or YIPF4 deletion is indicated. Proteins across all classes of transmembrane segments within the Golgi proteome were identified, although the majority of Golgi ‘Autophagy’ candidate proteins stabilized upon YIPF4 depletion contain a single transmembrane segment. Among the Golgi-associated proteins that are stabilized in FIP200^-/-^ cells, 5 out of 23 were stabilized (YIPF4^-/-^ log_2_(-AA/UT) – WT log_2_(-AA/UT)> 0.2) by YIPF4 deletion (**Figure 6C**), while 30 out of 54 Golgi-membrane proteins were stabilized. Consistent with our observation in HEK293 cells, correlation plots of YIPF3^-/-^ or YIPF4^-/-^ and ATG7^-/-^ from HeLa cells also show selectivity for Golgi membrane proteins (**fig. S7B, Data S6**). Thus, both YIPF3 and YIPF4 act as selective Golgiphagy receptors in two different cell lines.

**Figure 6.**
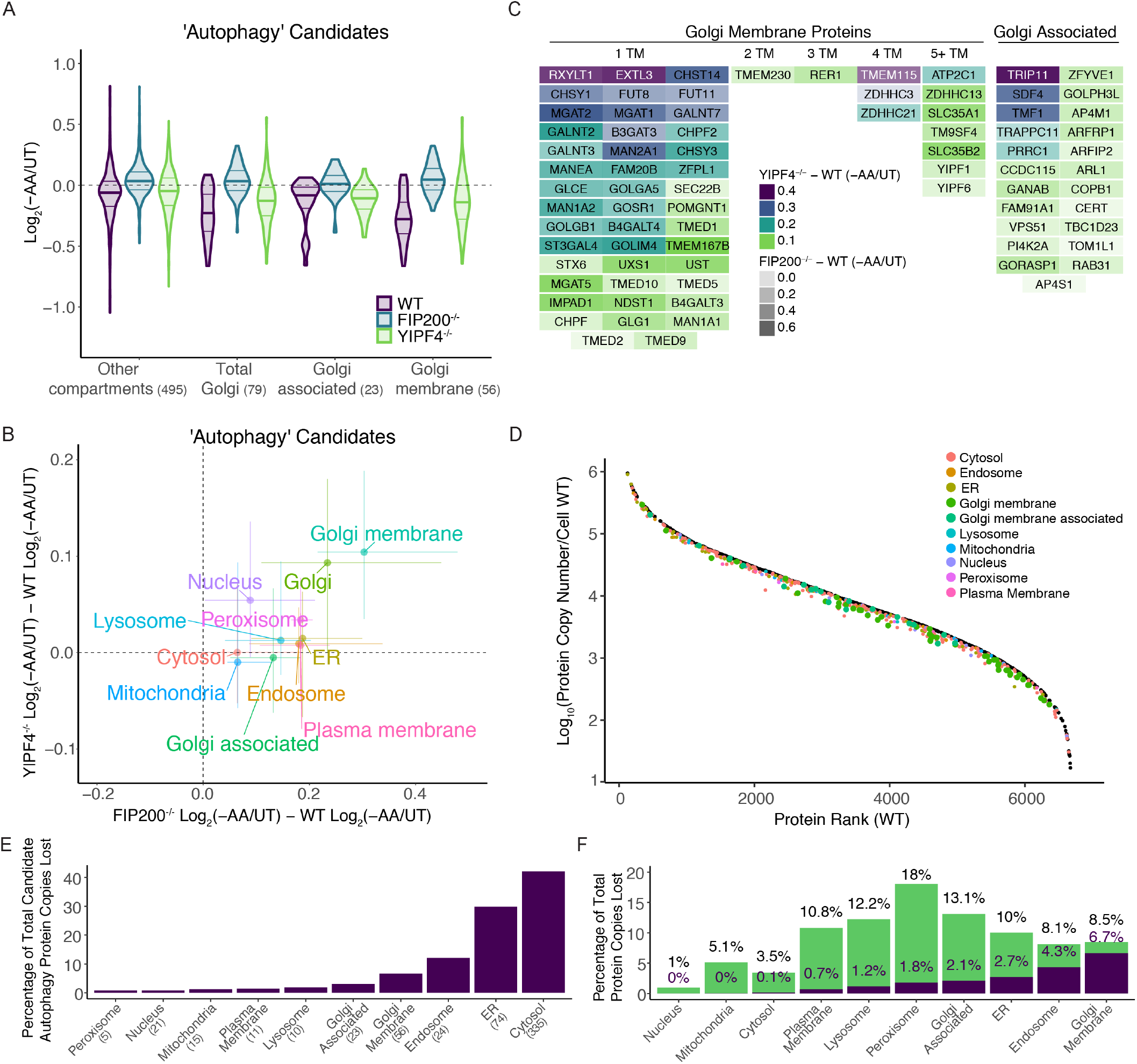
Protein census for autophagic protein degradation during nutrient stress. (**A**) Violin plots for total Golgi, Golgi-associated, Golgi-membrane, and other classes of proteins in WT, FIP200^-/-^, or YIPF4^-/-^ HEK293 cells in response to AA withdrawal (12 hours). Median value for the proteins within each group is indicated by the bold solid line. (**B**) Correlation plot of candidate ‘autophagy’ proteins for alterations in protein abundance for the indicated sub-cellular compartments during AA withdrawal for YIPF4^-/-^/WT cells (y-axis) versus FIP200^-/-^/WT cells (x-axis). Points are median of each distribution, and lines represent 25-75% quantile. (**C**) Classification of Golgi proteins that display YIPF4 or FIP200- dependent degradation in response to AA withdrawal (12 hours), with the number of trans-membrane segments for each membrane protein, as well as Golgi-associated proteins, shown. Grey density scale indicates the FIP200-dependence while the color scale indicates YIPF4-dependence. (**D**) TMT-scaled MS1 ranked plots. Protein copy number estimates for candidate ‘autophagy’ proteins in HEK293 cells (black) in rank order. Among the ‘autophagy’ candidate list, the number of protein copies after loss by autophagy during amino acid starvation for each compartment as determined using protein abundance fold changes (AA withdrawal – untreated). (**E**) Among the candidate autophagy proteins, percentage of total protein copy numbers lost via amino acid withdrawal. (**F**) Percentage of all protein copies lost from ‘autophagy’ candidate list (purple) or other mechanisms (green) by amino acid withdrawal for subcellular compartments (1.2829 x 10^6^ total).

Previous studies concluded that a soluble Ub-binding autophagy adaptor CALCOCO1 is involved in both Golgi and ER turnover during nutrient stress (*14, 15*). However, we found that the abundance of both YIPF3 and YIPF4, as measured by immunoblotting, was reduced in CALCOCO1^-/-^ HeLa cells in response to EBSS (18 hours) to a similar extent as seen in control cells, and this reduction was blocked by ATG7 deletion in the same experiment (**fig S8A**). Nutrient stress-dependent degradation of the ER-phagy receptor TEX264 was also not affected in CALCOCO1^-/-^ cells (**fig. S8A**). Proteomic analysis of cells lacking CALCOCO1 in response to EBSS revealed an extent of Golgi-membrane protein turnover comparable to control cells, while YIPF4^-/-^ cells in the same experiment displayed strong stabilization of Golgi-membrane proteins (**fig. S8B, Data S7**). Taken together, these data indicate that if CALCOCO1 is involved in Golgi-membrane turnover by autophagy, it functions downstream of YIPF4 or is involved in a distinct arm of the Golgiphagy response.

### Proteome census for autophagic cargo degradation with nutrient stress

A central question in the autophagy field concerns how cells determine substrates for autophagy in response to perturbations while maintaining cellular homeostasis. *a priori*, abundant cellular complexes might be considered as likely autophagy substrates to provide recycled amino acids without dramatically impacting cellular homeostasis. However, consistent with previous studies (*13*), we find that subunits of abundant cellular complexes such as the ribosome and proteasome do not reflect an autophagy dependent turnover profile that would be consistent with our candidate ‘autophagy’ proteins (**fig. S9A**). Likewise, while the cytosol accounts for ~59% of protein content in HeLa cells, other organelles such as Golgi and ER account for only a small fraction of the proteome (~0.8 and 4.4%, respectively) (*27*), yet their proteins are substantially enriched within the candidate ‘autophagy’ proteins identified here. This raises the question of how autophagic substrates scale with total protein abundance within the cell and across individual sub- cellular compartments.

To address this question, we merged estimates for absolute protein abundance and our quantitative proteome measurements upon starvation with the goal of providing a ‘proteome census’ for nutrient stress. Thus, we integrated protein copy number per cell with subcellular residency for protein molecules present in the cohort of candidate ‘autophagy’ proteins observed with 12 hours of AA withdrawal. First, we estimated protein copy number per cell using the Proteome Ruler method (*28*) extrapolated MS^1^ signal from relative TMT intensities (see **METHODS**) in untreated WT cells. We then inferred each protein’s loss in estimated absolute abundance based on the protein’s relative fold change in starvation conditions. Plots of autophagy-dependent protein copy number loss for each cellular compartment span ~5 orders of magnitude in abundance across ~6,800 proteins quantified, indicating that macroautophagy does not only degrade the most abundant cytosolic, ER, and Golgi proteins (**Figure 6D, fig. S9B**). In fact, the abundance ranks for ‘Autophagy’ candidate proteins is not significantly different from all other proteins. However, at the level of sub cellular compartments, organelles display differing degrees of selectivity (**fig. S9C**). Interestingly, cytosol ‘Autophagy’ candidates show a bias toward less abundant proteins, while ER and Endosome ‘Autophagy’ candidates are biased for more abundant proteins. In contrast, Golgi proteins in the ‘Autophagy’ candidate show no preference for more or less abundant Golgi proteins.

Based on our absolute abundance estimates, we calculated the total number of protein copies per cell that were degraded for candidate ‘autophagy’ proteins based on their primary subcellular compartment. The vast majority of protein copies degraded, as a percentage of the total candidate ‘autophagy’ molecules lost, are contributed by ER, endosome, Golgi, and cytosol, but unexpectedly, the number of protein molecules contributed by ER and Golgi rivals that of the cytosol (**Figure 6E, Data S8**). Given that proteasomal or ESCRT-dependent degradation and translational suppression also play a role in determining protein abundance during starvation (*12, 13*), we calculated the fractional contribution of protein abundance loss from each candidate ‘autophagy’ protein relative to the total abundance loss during starvation for individual compartments. ~75% of the reduction in protein abundance of Golgi membrane proteins could be attributed to the proteins that are prioritized for being turned over by autophagy, with endosomes and the ER also having a substantial amount of autophagy-based loss (**Figure 6F**). In contrast, only ~3% of the changes in the copy number of cytosolic proteins could be attributed to the abundance loss from the candidate ‘Autophagy’ proteins (**Figure 6F**). Analogous results were obtained when our data was mapped onto absolute abundance estimates previously reported in HEK293T cells (*28*) or derived from MS data measured by Data Independent Acquisition mass spectrometry (DIA) (**fig. S10A-J**), with absolute abundance estimates that correlated well with data herein (**fig S10A-J**). Thus, Golgi and ER represent major targets for autophagy in response to nutrient stress with a larger fraction of their individual proteomes being subjected to turnover than the cytosol, despite a much larger (>10-fold) copy number of cellular proteins being present within the cytosol (*27*).

## DISCUSSION

More than 30% of the cellular proteome is synthesized on the ER and trafficked through the Golgi, where proteins are modified and sent to other destinations. As such, these organelles are critical for cellular and organismal viability. Here we developed a prioritization strategy to categorize putative autophagy factors using complementary proteomic methods. This analysis allowed us to detail a pathway that mediates Golgi-membrane protein remodeling by autophagy in response to nutrient stress, thereby extending our understanding of mechanisms underlying the removal of membrane-bound organelles during starvation beyond well-studied ER-phagy pathways (*10*). We demonstrate that the Golgi resident protein YIPF4 is mobilized into vesicles in an autophagy-dependent process, is degraded by autophagy, and is required for autophagic degradation of a cohort of primarily Golgi-membrane proteins. Its heterodimeric partner YIPF3 likely functions together with YIPF4 in these processes, as we demonstrate that it also undergoes Golgiphagic flux, and its deletion led to stabilization of primarily Golgi-membrane proteins during nutrient stress. We have created a comprehensive <u>C</u>ellular <u>A</u>utophagy <u>R</u>egulation and <u>GO</u>lgiphagy (*CARGO*) web resource that allows exploration of all data generated reported here (**fig. S11A-C**).

In ER-phagy, LIR motifs in multiple transmembrane ER-phagy receptors are thought to nucleate autophagosome formation through interaction with FIP200/ULK1 and/or ATG8 proteins (*10*). Further work is required to understand the biochemical mechanisms underly coupling of YIPF3 and YIPF4 with the autophagy machinery, as well as whether Golgiphagy can be promoted via other types of signaling paradigms or is involved in Golgi quality control processes akin to those found with misfolded secretory proteins in ER-phagy (*10*). As with ER-phagy, it is possible that additional membrane-embedded Golgi proteins can promote selective Golgiphagy, possibly in diverse cell types or in response to distinct signals. Regardless of their specific activation signals or mechanism, our data establish a critical role of YIPF3 and YIPF4 in remodeling the Golgi proteome during nutrient stress.

By combining autophagy deficient cells, starvation, and consensus autophagy substrate profiling analysis using RMSE, we decoupled autophagic turnover from other mechanisms that decrease protein abundance during nutrient stress to identify a cohort of proteins that are degraded in an autophagy dependent manner. Although the RMSE approach may not capture every autophagy substrate (see METHODS), the prioritized collection of candidate ‘autophagy’ substrates nevertheless allowed us to create a ‘proteome census’ for nutrient stress by merging protein copy number estimates with subcellular compartment data. Historically, macroautophagy has been viewed as a non-specific process wherein bulk cytoplasmic proteins are captured for lysosomal degradation. Our results suggest an alternative model in which targeted degradation of ER and Golgi constitute major programs within a larger macroautophagy process (Figure 5H). Although ER and Golgi collectively accounting for ~6% of the protein copies per cell (*27*), a selective subset of their proteins within candidate ‘autophagy’ proteins account for ~50% of all protein copies lost (Figure 6E, F, fig. S10A-J), despite a much larger total copy number for cytosolic proteins within cells (~59% of cellular proteome) (*27*). These findings raise the question of whether autophagic degradation of cytosolic proteins during nutrient stress reflects bystander engulfment during selective autophagy of other organelles (principally ER and Golgi), or a program for selection/exclusion of specific cytosolic proteins (Figure 5I). Regardless, our data support idea that selective organelle-phagy represents a major component of macroautophagy. The preference for ER and Golgi could reflect the underlying mechanisms of membrane-templated autophagosome assembly that is frequently used to capture cargo via selective autophagy. Alternatively, while ER-derived phospholipids are used to make autophagosomal membranes via ATG2-dependent transport and are therefore recycled within the lysosome (*3*), it is possible that ER and Golgi have been evolutionarily programmed for autophagic targeting in order to provide additional classes of lipids present in these membranes for recycling in times of nutrient stress.

## ACKNOWLEDGMENTS

We thank members of the Harper lab for feedback. We acknowledge the Nikon Imaging Center (Harvard Medical School) for imaging assistance.

## Funding

This work was supported by Aligning Science Across Parkinson’s (ASAP) (JWH.).

NIH R01 NS110395 (JWH)

NIH R01 AG011085 (JWH)

NIH R01GM132129 (JAP)

Merck-Helen Hay Whitney Foundation (KLH)

Michael J Fox Foundation administers the grant ASAP-000282 on behalf of ASAP and itself.

For the purpose of open access, the author has applied for a CC-BY public copyright license to the Author Accepted Manuscript (AAM) version arising from this submission.

## Author Contributions

Conceptualization: SS, KLH, JWH

Investigation: SS, KLH, IRS, JCP, JAP

Analysis: KLH, IRS, SS

CARGO website creation: IRS

Visualization: KLH, IRS

Writing—original draft: KLH, JWH

Writing—reviewing and editing: KLH, SS, IRS, JCP, JAP, JWH

## Data and materials availability

All the mass spectrometry proteomics data (155.RAW files) have been deposited to the ProteomeXchange Consortium via the PRIDE repository (http://www.proteomexchange.org/): (Project Accession: PXD038358). All analyzed proteomic data are in Data S1, S2, S4, S5, S6, S7, and S8.

## Code and Software Availability

Code and data analysis to generate paper figures can be found on GitHub at https://github.com/harperlaboratory/Golgiphagy.git. All data and data figures can be explored using CARGO (Cellular Autophagy Regulation and GOlgiphagy). CARGO is a ShinyApp interface generated in R and RStudio that can be accessed at https://harperlab.connect.hms.harvard.edu/CARGO_CellularAutophagyRegulationandGOlgiphagy/.

## MATERIALS AND METHODS

### Reagents

#### Antibodies

ATG7 (Cell Signaling Technology, 8558S), LC3B (MBL international, M186-3), ULK1 (Cell Signaling Technology 8054), Phospho-ULK1 (ser757) (Cell Signaling Technology 14202), 4E-BP1 (Cell Signaling Technology 9644), Phospho-4E-BP1 (Thr37/46) (Cell Signaling Technology 2855), TEX264 (Sigma, HPA017739), Tubulin (Abcam, ab7291), 4EBP1 (Cell Signal Technology, 9644), YIPF3 (Invitrogen PA566621), YIPF4 (Sino Biological 202844-T46), HSP90 (Proteintech 60318), CALCOCO1 (Abclonal A7987), LAMP1 (Cell Signaling Technology 9091), GOLGB1/ Giantin (abcam ab37266), GOLGA2 (Proteintech 11308), PCNA (Santa Cruz PC10), IRDye 800CW Goat anti-Rabbit IgG H+L (LI-COR, 925-32211), IRDye 680 RD Goat anti-Mouse IgG H+L (LI-COR, 926-680),

#### Chemicals, Peptides, and Recombinant Proteins

FluoroBrite DMEM (Thermo Fisher Scientific A, 1896701), Benzonase Nuclease HC (Millipore, 71205-3), Urea (Sigma, Cat#U5378), SDS (Sodium Dodecyl Sulfate) (Bio-Rad, Cat#1610302), Dulbecco’s MEM (DMEM), high glucose, pyruvate (Gibco / Invitrogen, 11995), Dulbecco’s MEM (DMEM), Low Glucose, w/o Amino Acids (US Biological, D9800-13), TCEP (Gold Biotechnology), Puromycin (Gold Biotechnology, P-600-100), Protease inhibitor cocktail (Sigma-Aldrich, P8340), PhosSTOP (Sigma-Aldrich, 4906845001), Trypsin (Promega, V511C), Lys-C (Wako Chemicals, 129-02541), EPPS (Sigma-Aldrich, Cat#E9502), 2-Chloroacetamide (Sigma-Aldrich, C0267), TMT 11plex Label Reagent (Thermo Fisher Scientific, Cat#90406 & #A34807), TMTpro 16plex Label Reagent (Thermo Fisher Scientific, Cat#A44520), Hydroxylamine solution (Sigma Cat#438227), Empore™ SPE Disks C18 (3M - Sigma-Aldrich Cat#66883-U), Sep-Pak C18 Cartridge (Waters Cat#WAT054960 and #WAT054925), SOLA HRP SPE Cartridge, 10 mg (Thermo Fisher Scientific, Cat#60109-001), High pH Reversed-Phase Peptide Fractionation Kit (Thermo Fisher Scientific, Cat#84868), Bio-Rad Protein Assay Dye Reagent Concentrate (Bio-Rad,#5000006), and EBSS (Sigma-Aldrich Cat#E3024).

### Cell lines

HEK293 (human embryonic kidney, fetus, ATCC CRL-1573, RRID: CVCL_0045), and HeLa (cervical carcinoma cell line CCL-2; RRID: CVCL_0030) cells were grown in Dulbecco’s modified Eagle’s medium (DMEM, high glucose and pyruvate) supplemented with 10% fetal calf serum and maintained in a 5% CO_2_ incubator at 37°C. Cells were maintained at <80% confluency throughout the course of experiments. HeLa cells lacking MAP1LC3B or GABARAPL2 were from a previous study (*17*).

### Nutrient starvation experiments

Cells were plated in 6-well, 10cm or 15cm dishes the night before nutrient stress. DMEM was removed and cells were washed 3 times with DPBS followed by resuspending cells in EBSS or DMEM lacking amino acids prepared according to (*5*). For whole cell proteomics experiments, cells were resuspended in EBSS or media lacking amino acids as described in (*7*) for 12-18. For APEX2 proximity labeling and imaging experiments, cells were resuspended in EBSS+ BafA for 3-4 hours in the presence or absence of indicated inhibitors.

### CRISPR-Cas9 gene editing

*YIPF4, FIP200, ATG7* knock-out in HEK293, and *ATG7, YIPF4, CALCOCO1* knock-out in HeLa cell lines were carried out by plasmid-based transfection of Cas9/gRNA using pX459 plasmid as described (*29*). The following gRNAs, designed using the CHOPCHOP website (http://chopchop.cbu.uib.no/), were used: YIPF4: 5’ ATCTCGCGGCGACTCCCAAC / CGGCCTATGCCCCCACTAAC 3’; FIP200: 5’ ACTACGATTGACACTAAAGA 3’; ATG7 HEK293: 5’ ATCCAAGGCACTACTAAAAG 3’; CALCOCO1: 5’ AAGTTGACTCCACCACGGGA / CTAAGCCGGGCACCATCCCG 3’. Puromycin selection was followed 24-48 hours after the transfection. Cells were given a day to recover from puromycin selection and then single cells were sorted into a 96-well plate using fluorescence-activated cell sorting (FACS) on the SONY SH800S sorter. Individual clones were screened for deletion of the relevant gene by immunoblotting cell extracts with antibodies specific for the designed gene product. For amino-terminal tagging of the YIPF4 locus, the gRNA 5’ TCGCCGCGAGATGCAGCCTC 3’ was cloned into pX459 and co-transfected with a repair template containing an mNEON Green cassette flanked by homology arms (pSMART-mNEON-YIPF4) into HEK293 and HEK293 FIP200^-/-^ cells using lipofectamine 3000. After 7 days, a population of cells for both genotypes was sorted for the same level of mNEON Green signal.

### Cell lysis and immunoblotting assay

Cells were cultured in the presence of the corresponding stress to 60-80% confluency in 6-well plates, 10 cm or 15 cm dishes. After removing the media, the cells were washed with DPBS three times. To lyse cell urea buffer (8M urea, 50 mM TRIS 7.5, 150 mM NaCl, containing mammalian protease inhibitor cocktail (Sigma), Phos-STOP, and 20 unit/ml Benzonase (Millipore)) was added directly onto the cells. Cell lysates were collected by cell scrapers and sonicated on ice for 10 seconds at level 5, and lysates were cleared by centrifugation (15000 rpm, 10 min at 4 °C). The concentration of the supernatant was measured by BCA assay. For immunoblotting, the whole cell lysate was denatured by the addition of LDS sample buffer supplemented with 100 mM DTT, followed by boiling at 95°C for 5 minutes. 10-20 μg of each lysate was loaded onto the 4-20% Tris-Glycine gel (Thermo Fisher Scientific) or 4-12% NuPAGE Bis-Tris gel (Thermo Fisher Scientific), followed by SDS-PAGE with Tris-Glycine SDS running buffer (Thermo Fisher Scientific) or MOPS SDS running buffer (Thermo Fisher Scientific), respectively. *For Chemiluminescence westerns:* The proteins were electro-transferred to PVDF membranes (0.45 μm, Millipore), and then the total protein was stained using Ponceau (Thermo Fisher Scientific). The membrane was then blocked with 5% non-fat milk (r.t., 60 min) incubated with the indicated primary antibodies (4°C, overnight), washed three times with TBST (total 30 min), and further incubated either with HRP conjugated anti-Rabbit and anti-mouse secondaries at (1:5,000) for 1 hour. After thorough wash with TBST for 30 min membranes were treated with Lightning™ Plus Chemiluminescence Reagent (PerkinElmer, NEL104001EA) after mixing the Enhanced Luminol Reagent and the Oxidizing Reagent 1:1. Mixed Chemiluminescence Reagent was added to blot and rocked gently for 1 minute and imaged using BioRad ChemiDoc Imaging System. *For LI-COR westerns:* The proteins were electro-transferred to nitrocellulose membranes and then the total protein was stained using Ponceau (Thermo Fisher Scientific). The membrane was then blocked with LI-COR blocking buffer at room temperature for 1 hour. Then membranes were incubated with the indicated primary antibodies (4°C, overnight), washed three times with TBST (total 30 min), and further incubated either with fluorescent IRDye 680RD Goat anti-Mouse IgG H+L, or IRDye 800CW Goat anti-Rabbit IgG H+L secondary antibody at (1:10,000) at room temperature for 1 hour. After thorough wash with TBST for 30 min, near infrared signal was detected using OdysseyCLx imager and quantified using ImageStudioLite (LI-COR).

### Flow Cytometry for Keima analysis

Corresponding cells were plated onto 96-well plates one day prior to the nutrient stress. The cells were washed twice with PBS and resuspended in DMEM or EBSS to start 16-hour starvation. After starvation, cells were treated with trypsin and quenched with Phenol red free-DMEM. Cells were filtered and analyzed by flow-cytometry (Attune NxT, Thermo Fisher) using the high throughput autosampler (CyKick). The data was processed by FlowJo software and plotted using GraphPad Prism.

### Confocal Microscopy

Cells were plated onto 18 or 22 mm-glass coverslips (No. 1.5, 22×22 mm glass diameter, VWR 48366-227) the day before nutrient stress. DMEM was removed and cells were washed three times with DPBS, followed by resuspension in EBSS + the appropriate inhibitor(s) (SAR405, BafA, TAK243). After starvation treatment, cells were fixed using 4% PFA followed by permeabilization with 0.5% Triton-X100. Cells were blocked in 3% BSA for 30 minutes, followed by incubation in primary antibodies for 1 hour at room temperature. Cells were washed 3 times with DPBS + 0.02% tween-20, followed by incubation in secondary (alexafluor conjugated secondary antibodies) for 1 hour at room temperature. Coverslips were then washed 3 times with DPBS + 0.02% tween-20 and mounted onto glass slides using mounting media (Vectashield H-1000) and sealed with nail polish. The cells were imaged using a Yokogawa CSU-W1 spinning disk confocal on a Nikon Ti motorized microscope equipped with a Nikon Plan Apo 100x/1.40 N.A objective lens, and Hamamatsu ORCA-Fusion BT CMOS camera. For the analysis, the equal gamma, brightness, and contrast were applied for each image using FiJi software. For quantification, at least 3 separate images were quantified for the number of mNEON puncta and nuclei.

### Proteomics Workflow

#### TMT total proteome sample preparation

Cells were cultured to 70% confluency and washed with PBS three times. Cells were lysed by in UREA denaturing buffer (8M Urea, 150mM NaCl, 50mM EPPS pH8.0, containing mammalian protease inhibitor cocktail (Sigma), and Phos-STOP) Cell lysates were collected by cell scrapers and sonicated on ice for 10 seconds at level 5, and resultant extracts were clarified by centrifugation for 10 minutes at 15,000xg at 4 °C. Lysates were quantified by BCA and ~50 μg of protein was reduced with TCEP (10mM final concentration for 30 min) and alkylated with Chloroacetamide (20mM final concentration) for 30 minutes. Proteins were chloroform-methanol precipitated using the protocol in SL-TMT protocol (*30*), reconstituted in 200 mM EPPS (pH 8.5), digested by Lys-C for 2 hours at 37 degrees (1:200 w:w LysC:protein) and then by trypsin overnight at 37°C (1:100 w:w trypsin:protein). ~25μg of protein was labeled with 62.5 μg of TMT or TMTpro for 120 min at room temperature. After labeling efficiency check, samples were quenched with hydroxylamine solution at ~0.3% final (w. in water), pooled, and desalted C18 solid-phase extraction (SPE) (SepPak, Waters). Pooled samples were offline fractionated with basic reverse phase liquid chromatography (bRP-LC) into a 96-well plate and combined for a total of 24 fractions (*31*) before desalting using a C18 StageTip (packed with Empore C18; 3M Corporation), and subsequent LC–MS/MS analysis.

#### DIA total proteome sample preparation

HEK293 cells (with or without amino acid withdrawal treatment) were cultured to ~70% confluency, washed twice with chilled PBS, and harvested by cell scraping in PBS. Following centrifugation at 4°C, cell pellets were lysed in a denaturation buffer (8M Urea, 150mM NaCl, 50mM EPPS pH8.0, containing mammalian protease inhibitor cocktail (Sigma), and Phos-STOP) by sonication (three times at level 5 for 5 seconds, with rest on ice). Cell extracts were clarified by centrifugation for 10 minutes at 15,000xg at 4°C. Lysates were quantified by BCA and protein was reduced with TCEP (5 mM final concentration for 30 min), alkylated with IAA (10 mM final concentration) in the dark for 30 minutes, and quenched with DTT (5 mM final concentration) for 30 minutes. 100 ug of protein was methanol-chloroform precipitated using the protocol in SL-TMT protocol (*30*), reconstituted in 100 mM EPPS (pH 8.5 at 1 mg/mL), digested by Lys-C for 2 hours at 37°C (1:100 w:w LysC:protein) and then by trypsin overnight at 37°C (1:100 w:w trypsin:protein). 30 ug of protein digests were acidified with formic acid to pH ~3-3.5, desalted using a C18 StageTip (packed 200ul pipette tip with Empore C18; 3M Corporation), and subjected to data independent acquisition (DIA) LC-MS/MS analysis.

#### Sample preparation for Mass Spectrometry-APEX2 Proteomics

For APEX2 proteomics, cells expressing various APEX2 fusions were processed as in (*18*).To induce proximity labeling in live cells, cells were incubated with 500 μM biotin phenol (LS-3500.0250, Iris Biotech) for 1 hr and treated with 1 mM H2O2 for 1 min, and the reaction was quenched with three washes of 1× PBS supplemented with 5 mM Trolox, 10 mM sodium ascorbate and 10 mM sodium azide. Cells were then harvested and lysed in radioimmunoprecipitation assay (RIPA) buffer. To enrich biotinylated proteins, ~2mg of cleared lysates was subjected to affinity purification by incubating with the streptavidin-coated agarose beads (catalog no. 88817, Pierce) for 1.5 hours at room temperature. Beads were subsequently washed twice with RIPA buffer, once with 1 M KCl, once with 0.1 M NaCO_3_, once with PBS and once with water. For proteomics, biotinylated protein bound to the beads were reduced using TCEP (10mM final concentration) in EPPS buffer at room temperature for 30 minutes. After reduction, samples were alkylated with the addition of Chloracetamide (20mM final concentration) for 20 minutes. Beads were washed three times with water. Proteins bound to beads were then digested with LysC (0.5μl) in 100ul of 0.1M EPPS (pH 8.5) for 2 hours at 37°C, followed by trypsin overnight at 37°C (1 μl). To quantify the relative abundance of individual protein across different samples, each digest was labeled with 62.5 μg of TMT11 or TMT16pro reagents for 2 hours at room temperature (Thermo Fisher Scientific), mixed, and desalted with a C18 StageTip (packed with Empore C18; 3M Corporation) before SPS-MS^3^ analysis on an Orbitrap Fusion Lumos Tribrid Mass Spectometer (Thermo Fisher Scientific) coupled to a Proxeon EASY-nLC1200 liquid chromatography (LC) pump (Thermo Fisher Scientific). Peptides were separated on a 100 μm inner diameter microcapillary column packed with ~35 cm of Accucore150 resin (2.6 μm, 150 Å, ThermoFisher Scientific, San Jose, CA) with a gradient consisting of 5%–21% (ACN, 0.1% FA) over a total 150 min run at ~500 nL/min (*32*). Details of instrument parameters for each experiment are provided below.

#### TMT Data acquisition

Samples were analyzed on Orbitrap Fusion Lumos Tribrid Mass Spectrometer coupled to a Proxeon EASY-nLC 1200 pump (ThermoFisher Scientific). Peptides were separated on a 35 cm column packed using a 95 to 110 min gradient. MS^1^ data were collected using the Orbitrap (120,000 resolution). MS^2^ scans were performed in the ion trap with CID fragmentation (isolation window 0.7 Da; rapid scan; NCE 35%). Each analysis used the Multi-Notch MS^3^-based TMT method (*32*), to reduce ion interference compared to MS^2^ quantification, combined in some instance with newly implemented Real Time Search analysis (*33, 34*), and with the FAIMS Pro Interface (using previously optimized 3 CV parameters (−40, −60, −80) for TMT multiplexed samples (*35*)). MS^3^ scans were collected in the Orbitrap using a resolution of 50,000, NCE of 65 (TMT) or 45 (TMTpro). The closeout was set at two peptides per protein per fraction, so that MS^3^ scans were no longer collected for proteins having two peptide-spectrum matches (PSMs) that passed quality filters.

#### DIA data acquisition

Samples were analyzed on an Orbitrap Exploris 480 Mass Spectrometer coupled to a Proxeon EASY-nLC pump 1000 (ThermoFisher Scientific). Peptides were separated on a 15 cm column packed with Accucore150 resin (150 Å, 2.6mm C18 beads Thermo Fisher Scientific, San Jose, CA) using an 80 min acetonitrile gradient. MS^1^ data were collected using the Orbitrap (60,000 resolution, 350-1,050 m/z, 100% Normalized AGC, maxIT set to “auto”). DIA MS^2^ scans in the Orbitrap were performed overlapping 24 m/z windows for first duty cycle (390-1,014 m/z) and for second duty cycle (402-1,026 m/z) with 28% NCE, 30,000 resolution, for fixed 145-1,450 m/z range, 1,000% normalized AGC, and 54 ms maxIT MS^1^ survey scan was performed following each DIA MS/MS duty cycle.

#### TMT Data analysis

Mass spectra were converted to mzXML and monoisotopic peaks were reassigned with Monocole (*36*) and then database searched using a Sequest-based (*37, 38*) or Sequest-HT using Proteome Discoverer (v2.3.0.420 – Thermo Fisher Scientific). Database searching included all canonical entries from the Human reference proteome database (UniProt Swiss-Prot – 2019-01; https://ftp.uniprot.org/pub/databases/uniprot/previous_major_releases/release-2019_01/) and sequences of common contaminant proteins. Searches were performed using a 20 ppm precursor ion tolerance, and a 0.6 Da product ion tolerance for ion trap MS/MS were used. TMT tags on lysine residues and peptide N termini (+229.163 Da for Amino-TMT or +304.207 Da for TMTpro) and carbamidomethylation of cysteine residues (+57.021 Da) were set as static modifications, while oxidation of methionine residues (+15.995 Da) was set as a variable modification. PSMs were filtered to a 2% false discovery rate (FDR) using linear discriminant analysis as described previously (*37*) using the Picked FDR method (*39*), proteins were filtered to the target 2% FDR level. For reporter ion quantification, a 0.003 Da window around the theoretical m/z of each reporter ion was scanned, and the most intense m/z was used. Peptides were filtered to include only those peptides with >200 summed signal-to-noise ratio across all TMT channels. An isolation purity of at least 0.5 (50%) in the MS1 isolation window was used for samples analyzed without online real-time searching. For each protein, the filtered peptide-spectral match TMT or TMTpro raw intensities were summed and log_2_ normalized to create protein quantification values (weighted average). Using protein TMT quantifications, TMT channels were normalized to the summed (protein abundance experiments) (*40*) or median (proximity labeling experiments) (*16*) TMT intensities for each TMT channel (adjusted to the median of the TMT channels summarization).

#### DIA data analysis

Mass spectra were converted to mzML using msconvert (*41*) with demultiplexing (Overlap Only at 10 ppm mass error). mzML files were processed with DIA-NN (*42*) using UniProt entries (UP000005640 [9606]). For DIA-NN, we used the following parameters: trypsin specificity ([RK]/P), N-term methionine excision enabled, fixed modification of carbamidomethylation on cysteines, in library-free mode, deep learning-based spectra and RTs enabled, MBR enabled, precursor FDR 1% filter, and quantification with Robust LC (high precision). Using the report.pg_matrix.tsv output from DIA-NN, we calculated the mean intensity across replicates for untreated and amino acid withdraw treatment conditions (n=4 each) based on replicate intensities (observed in at least two biological replicates) which were used to estimate a protein copy number per cell using the Proteome Ruler method (*28*).

### Statistics

Normalized log_2_ protein reporter ion intensities were compared using a Student’s t-test and resultant p-values were corrected using the Benjamini-Hochberg adjustment (Benjamini and Hochberg 1995). Volcano plots and other data visualizations were generated in R using resulting q-values and mean fold changes. Annotations for subcellular lists were derived from (*27*) and designations from (*43*). Additional cytosol protein and Golgi transmembrane number annotations were derived from Uniprot. GO annotations from Uniprot were appended to MS data to perform Fisher’s Exact tests to identify GO enrichment terms (corrected by Benjamini-Hochberg adjustment). Proteome ruler values were estimated using previously described methods (*28, 44*). The proportional contribution of the untreated WT TMT channels to the MS1 precursor are (TMT^WT/UT^ / TMT^All^ * MS1^Area^) was summed to the protein-level for its constituent peptides. Resultant protein values were then used to calculate a TMT-based proteome ruler protein absolute abundance estimate. For imaging quantification, a Mann-Whitney p-value was calculated using GraphPad Prism9. P-values <0.05 were considered significant unless otherwise noted. Compartment protein copy number rank tests were performed using a Wilcoxon test to calculate p-value. All data figures were generated in Adobe Illustrator, using R (4.1.3), Rstudio IDE(2021.09.3 Build 396, Posit), and GraphPad Prism9.

### RMSE calculation

To generate our candidate ‘autophagy’ protein list, we used known autophagy fluxers in autophagy proficient (WT) or deficient (ATG7^-/-^ or FIP200^-/-^) cells. For each known autophagy fluxer, the condition median z-score was used. From these protein condition medians, we took the median value across the known subset of proteins to estimate a condition median to build a consensus profile. Using the consensus profile median values for known autophagy proteins as predicted, we then calculated the RMSE for each protein in the data sets.

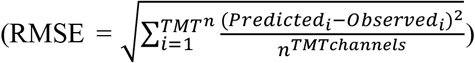. By calculating the RMSE for every quantified protein, we generated a group of candidate ‘autophagy’ proteins in two distinct starvation conditions based on the top 10% of proteins with the lowest RMSE across the datasets respectively. The 10% cutoff aligns well with the right most tail of the density plot for the known autophagy fluxers and the Top30 autophagy factors from **Figure 2**. While the resulting ‘autophagy’ candidate list provides a defined collection of autophagy substrates, the RMSE calculation averages the error across a protein’s abundance profile, potentially enabling some proteins that vary from the consensus profile in a single condition to make the candidate list. Also, some autophagy substrates with high replicate variance in abundance may not make the cutoff required despite largely following the known autophagy fluxer consensus profile.

### Prioritization of ‘autophagy’ cargo

To prioritize the top candidate autophagy cargo, we ranked proteins based on their starvation and autophagy turnover (**Figure 1**) and proximity to ATG8 machinery (**Figure 2**). To calculate a rank for starvation and autophagy dependent turnover, we determined the priority value based on the lesser of either the absolute value of the WT log_2_ fold change in protein abundance from EBSS/Untreated for log_2_(EBSS/UT) ≤ 0 or the ATG7^-/-^ log_2_(EBSS/UT) – WT log_2_(EBSS/UT) for changes ≥ 0 (when both criteria are met). Proteins that did not meet both criteria were assigned a 0 priority. The priority values were then arranged in descending order and proteins were scaled ranked (Protein Rank/Number of total proteins in the experiment). Scaled ranks were calculated for HeLa and HEK293 data separately and the minimum scaled rank found in at least one of the datasets was used. Proteins were reordered based on priority and scaled ranked combining the two datasets to summarize **Figure 1** findings. For ATG8 proximity ranks, we determined a priority value based on the lesser of either the log_2_ fold change in protein abundance from WT EBSS+Baf/Untreated for log_2_(EBSS+BafA/UT) ≥ 0 or the absolute value of the ATG8 LSD mutant log_2_(EBSS+BafA/UT) – WT log_2_(EBSS+BafA/UT) for changes ≤ 0 (only when both criteria are met). As above, proteins that did not meet both criteria were assigned a 0 priority. Using the priority values, scaled ranks were calculated for the APEX2-GABRAPL2 and APEX2-MAP1LC3B experiments separately, where the minimum scaled rank found in at least one of the experiments was used. Proteins were reordered based on priority and scaled ranked combining the two datasets to summarize **Figure 2** findings. To prioritize to candidates that were both turning over in an autophagy and starvation dependent manner and increased association with ATG8 during starvation, we summed **Figure 1** and **Figure 2** scaled ranks to generate a summed rank value that we sorted by in ascending order to generate our final ranked list of candidates. To be a candidate in the final ranked list the protein must have been identified in at least one experiment from **Figure 1** experiments (HeLa or HEK293) and one experiment from **Figure 2** experiments (APEX2-GABARAPL2 and APEX2- MAP1LC3B). LIR motifs were matched from iLIR Autophagy Database (http://repeat.biol.ucy.ac.cy/iLIR/) (*19*). Known autophagy proteins were derived from (*10*).

### Reproducibility

All experiments were repeated at least three times unless otherwise indicated.

### Data reporting

No statistical methods were used to predetermine sample size. The experiments were not randomized, and the investigators were not blinded to allocation during experiments and outcome assessment.

## SUPPLEMENTARY FIGURES

**fig. S1.**
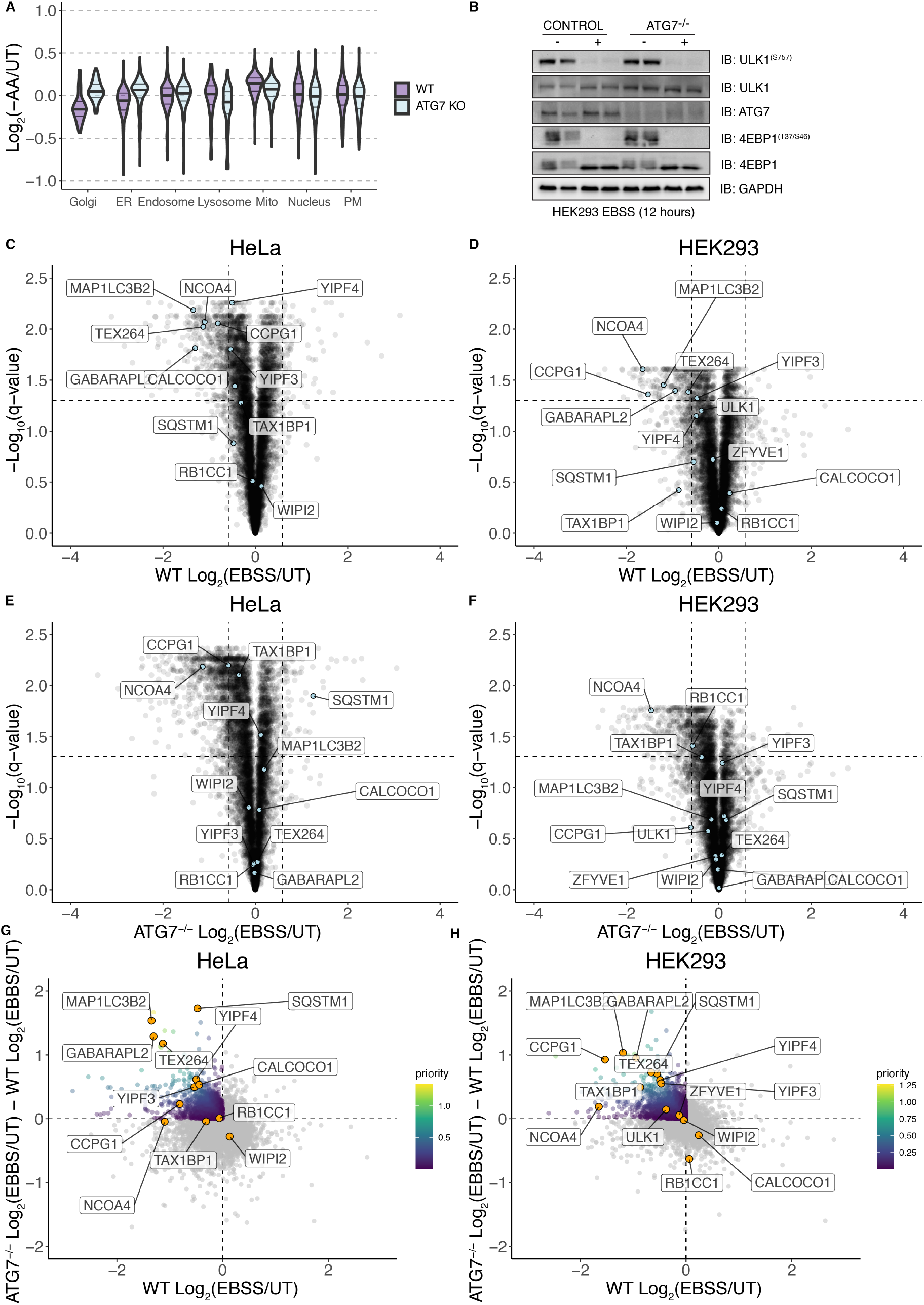
**(A)** Volcano plots for the indicated organelles in HEK293T cells with or without ATG7 in response to amino acid withdrawal (10 hours). Data are from our prior studies (*5, 13*). (**B**) Western blot showing markers of starvation (ULK1, 4EBP dephosphorylation) and ATG7 in WT and ATG7^-/-^ HEK293 cells grown in EBSS for 12 hours. (**C-F**) Volcano plots [WT Log_2_(12 hours EBSS/UT) versus -Log_10_(q-value)] for HeLa (panel C) or HEK293 (panel D) or analogous plots for ATG7^-/-^ HeLa (panel E) or HEK293 (panel F) cells. (**G,H**) Plots of ATG7^-/-^Log_2_(EBSS/UT) - WT Log_2_(EBSS/UT) versus WT Log_2_(EBSS/UT) for HeLa cells (panel E) and ATG7^-/-^ Log_2_(EBSS/UT) – WT Log_2_(EBSS/UT) versus WT Log_2_(EBSS/UT) for HEK293 cells (panel D) where priority for individual proteins is scaled based on the color code inset.

**fig. S2.**
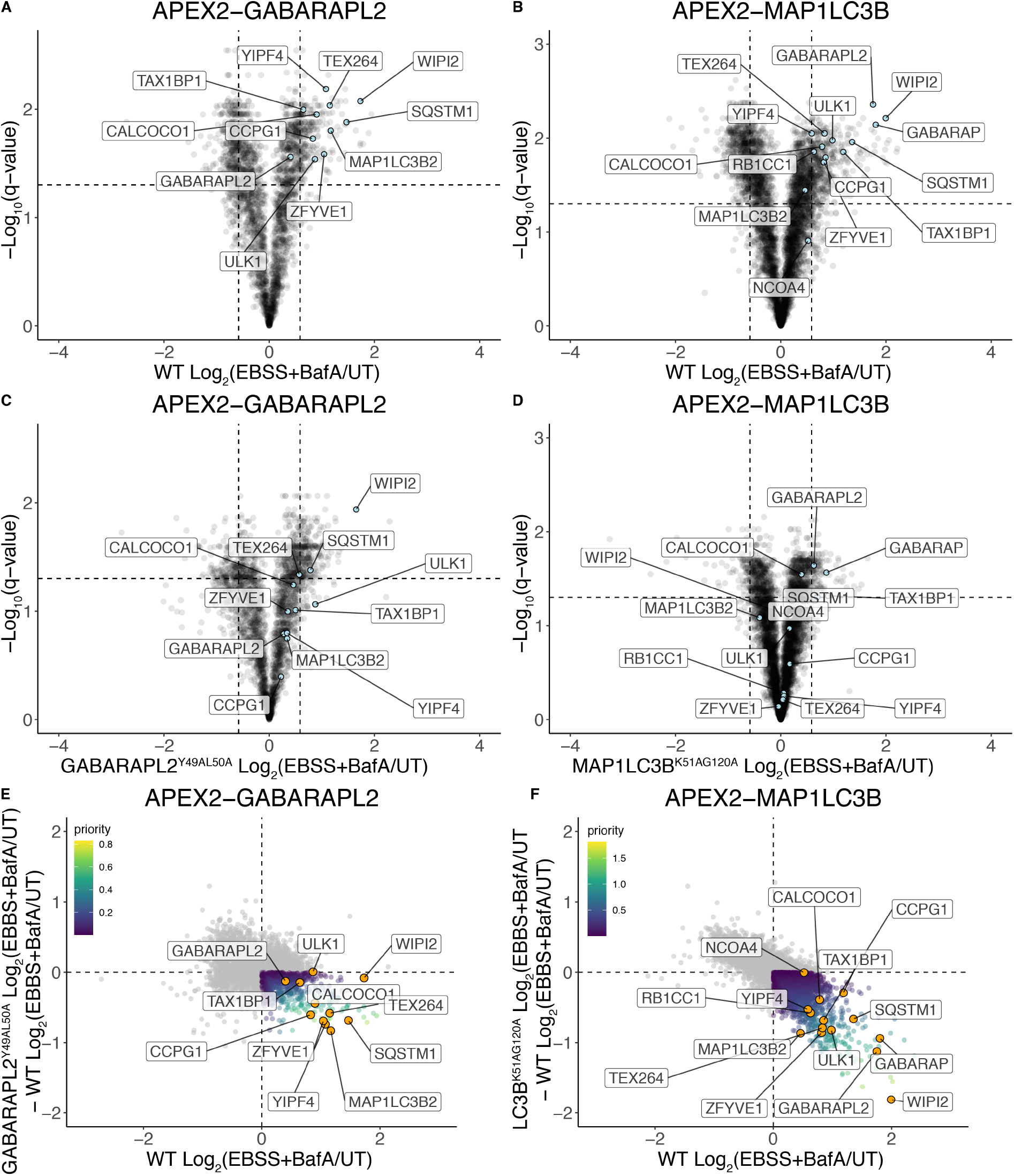
(**A-D**) Volcano plots [WT Log_2_(4 hours EBSS+BafA/UT) versus -Log_10_(q-value)] for APEX2- GABARAPL2 (panel A) or APEX2-MAP1LC3B (panel B) or analogous plots for APEX2- GABARAPL2^Y49A/L50A^ (panel C) or APEX2-MAP1LC3B^K51A/G120A^ (panel D) in HeLa cells. (**E**, **F**) Plots of GABARAPL2^Y49A/L50A^ Log_2_(EBSS+BafA/UT) – WT Log_2_(EBSS+BafA/UT) versus WT Log_2_(EBSS+BafA/UT) (panel E) and MAP1LC3B^K51A/G120A^ Log_2_(EBSS+BafA/UT) – WT Log_2_(EBSS+BafA/UT) versus WT Log_2_(EBSS+BafA/UT) (panel F) where priority for individual proteins is scaled based on the color code inset.

**fig. S3.**
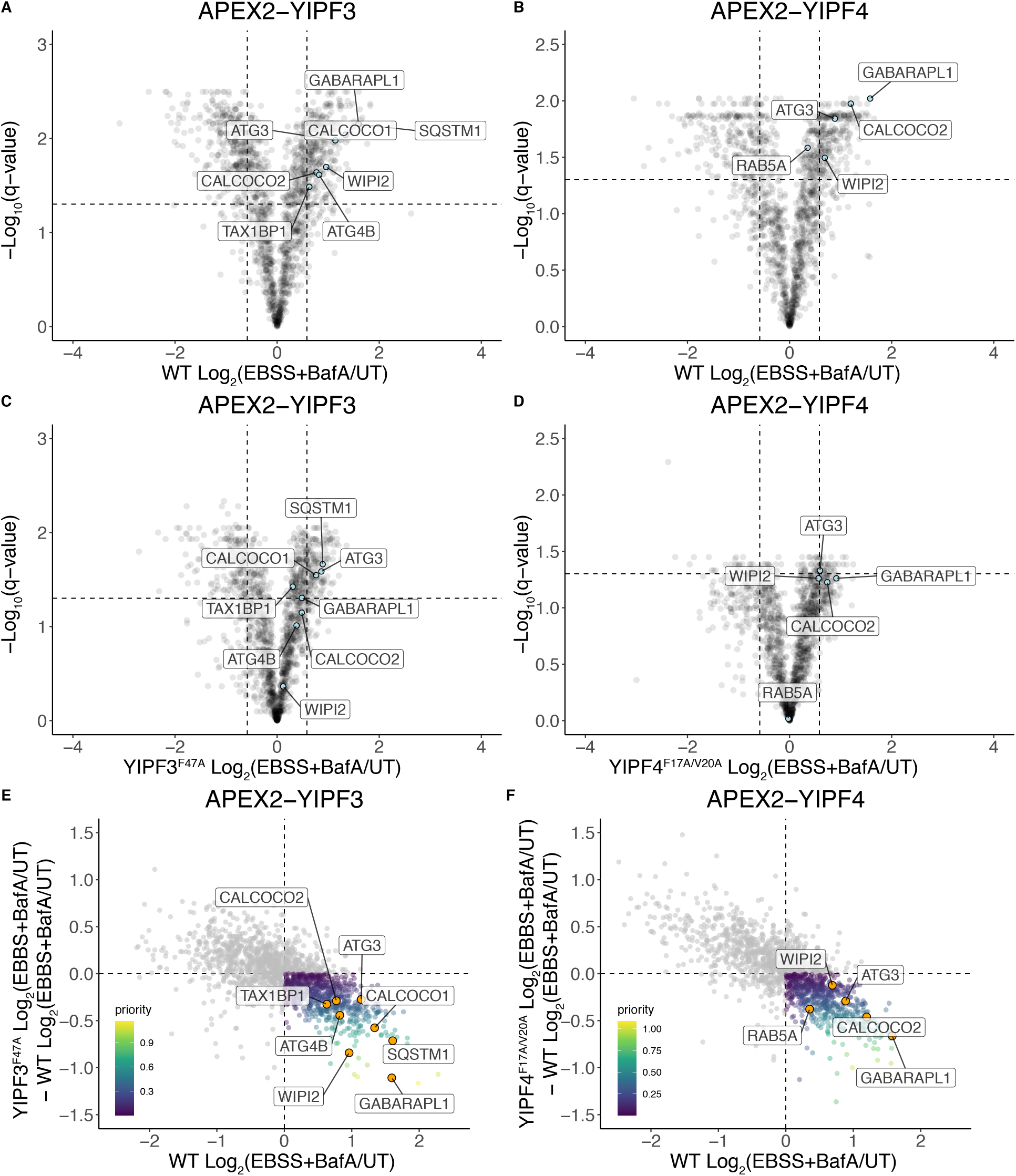
(**A-D**) Volcano plots [WT Log_2_(4 hours EBSS+BafA/UT) versus -Log_10_(q-value)] for APEX2- YIPF3 (panel A) or APEX2-YIPF4 (panel B) or analogous plots for APEX2-YIPF3^F47A^ (panel C) or APEX2-YIPF4^F17A/V20A^ (panel D) in HeLa cells. (**E, F**) Plots of APEX2-YIPF3^F47A^ Log_2_(EBSS+BafA/UT) WT Log_2_(EBSS+BafA/UT) versus WT Log_2_(EBSS+BafA/UT) (panel E) and APEX2-YIPF4^F17A/V20A^Log_2_(EBSS+BafA/UT) WT Log_2_(EBSS+BafA/UT) versus WT Log_2_(EBSS+BafA/UT) (panel F) where priority for individual proteins is scaled based on the color code inset.

**fig. S4.**
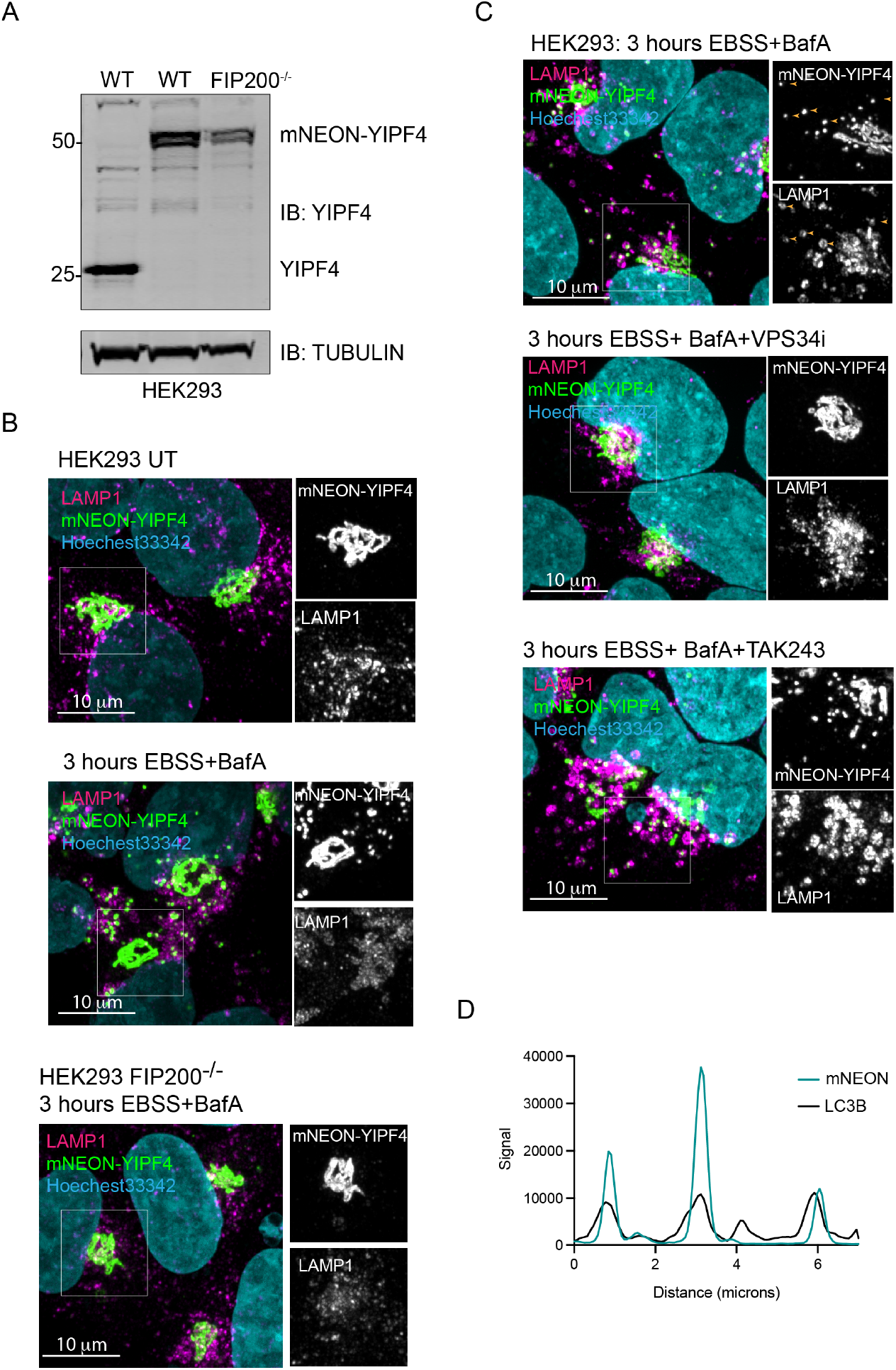
(**A**) Immunoblot showing mNEON-YIPF4 endogenous tagging results at a higher molecular weight indicative of the total fusion protein length (~50kDa). (**B**) HEK293 cells expressing endogenous YIPF4 tagged on its N-terminus with mNEON (green) imaged using confocal microscopy and co-stained with LAMP1 (magenta). Cells were either left untreated (top) or subjected to nutrient stress +BafA (3 hours) in wt or FIP200 ^-/-^ cells (middle and bottom) prior to imaging. Nuclei were labeled with Hoechst33342 dye (cyan). Scale bars 10 microns as indicated. (**C**) HEK293 cells expressing endogenous YIPF4 tagged on its N-terminus with mNEON (green) imaged using confocal microscopy and co-stained with LAMP1 (magenta). Cells were either left untreated (top) or subjected to nutrient stress +BafA and VPS34i (3 hours) (middle) or subjected to nutrient stress +BafA and an E1 inhibitor (TAK243) (3 hours) prior to imaging. Nuclei were labeled with Hoechst33342 dye (cyan). Scale bars 10 microns as indicated. (**D**) Line scan of HEK293 cells expressing endogenous YIPF4 tagged on its N-terminus with mNEON and MAP1LC3B show colocalization upon EBSS+BafA treatment for 3 hours.

**fig. S5.**
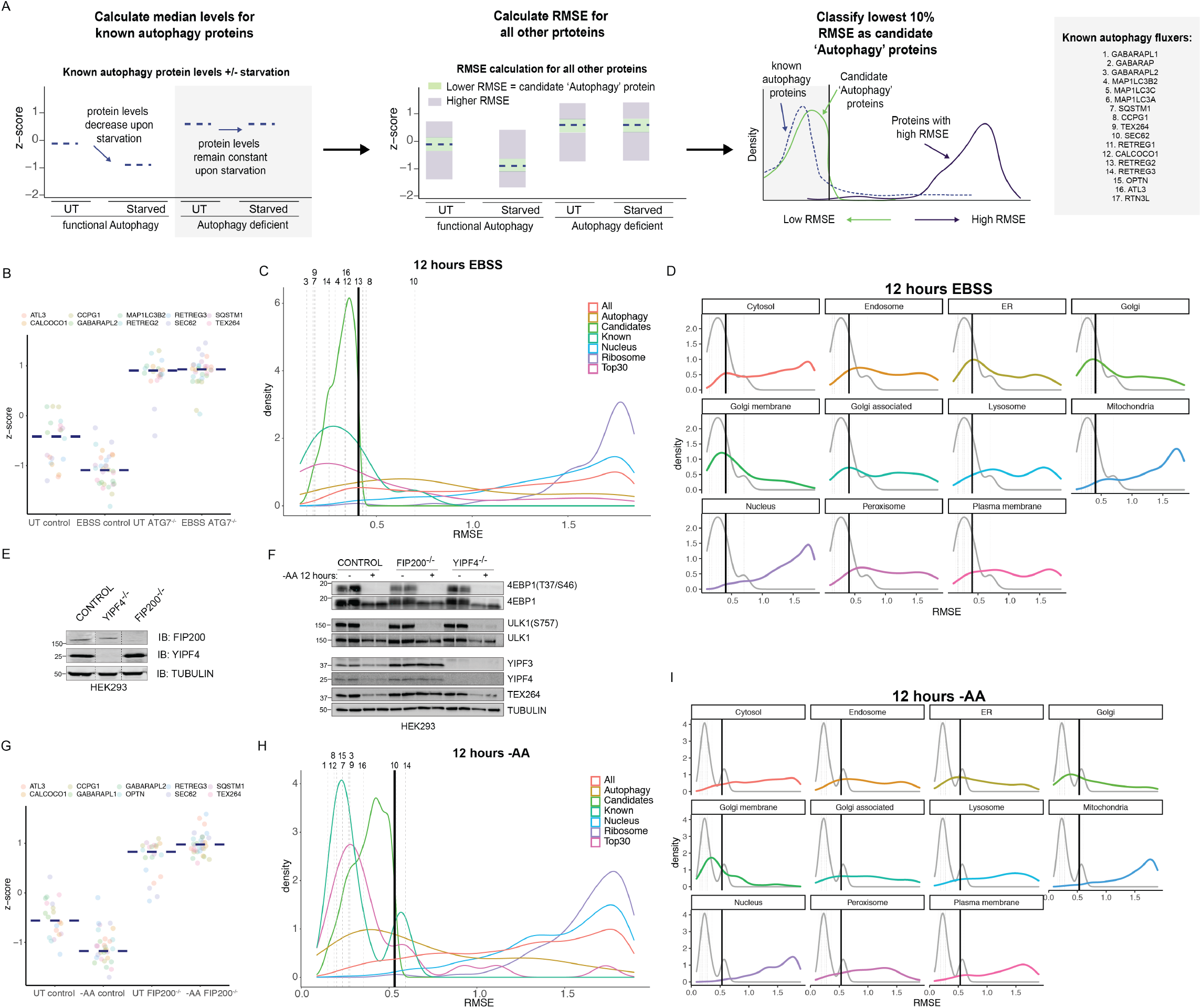
(**A**) Workflow for calculating RMSE of all proteins in HEK293 Control or ATG7^-/-^ cells treated with EBSS for 12 hours, and HEK293 control, FIP200^-/-^, and YIPF4^-/-^ cells treated with AA withdrawal for 12 hours. Known autophagy fluxers are shown on the right. (**B**) Dot plot of known autophagy fluxers in WT or ATG7^-/-^ HEK293 cells treated with EBSS for 12 hours. Navy dashed line represents median protein abundance. (**C**) RMSE plot for HEK293 cells treated with 12 hours of EBSS. Known autophagy proteins RMSE is shown. Top30 shows the top 30 proteins from the autophagy prioritization list generated from figures 1 and 2. Candidate ‘Autophagy’ proteins are shown along with all autophagy, nuclear, and ribosomal proteins. Grey vertical dashed lines represent each known autophagy fluxer quantified in the experiment, numbers represent the proteins according to panel A. (**D**) RMSE for each compartment shown from HEK293 EBSS experiment. (**E**) Immunoblot for control, FIP200^-/-^, and YIPF4^-/-^ cells with the indicated antibodies. (**F**) Immunoblot for HEK293 control, FIP200^-/-^ and YIPF4^-/-^ cells with or AA withdrawal for 12 hours in duplicate with the indicated antibodies. (**G**) Dot plot of known autophagy fluxers in WT or ATG7^-/-^ HEK293 cells treated with EBSS for 12 hours. Navy dashed line represents median protein abundance. (**H**) RMSE plot for HEK293 cells treated with 12 hours of AA withdrawal. Known autophagy proteins RMSE is shown. Top30 shows the top 30 proteins from the autophagy prioritization list generated from figures 1 and 2. Candidate ‘Autophagy’ proteins are shown along with all autophagy, nuclear, and ribosomal proteins. (**I**) RMSE for each compartment shown from HEK293 AA withdrawal experiment.

**fig. S6.**
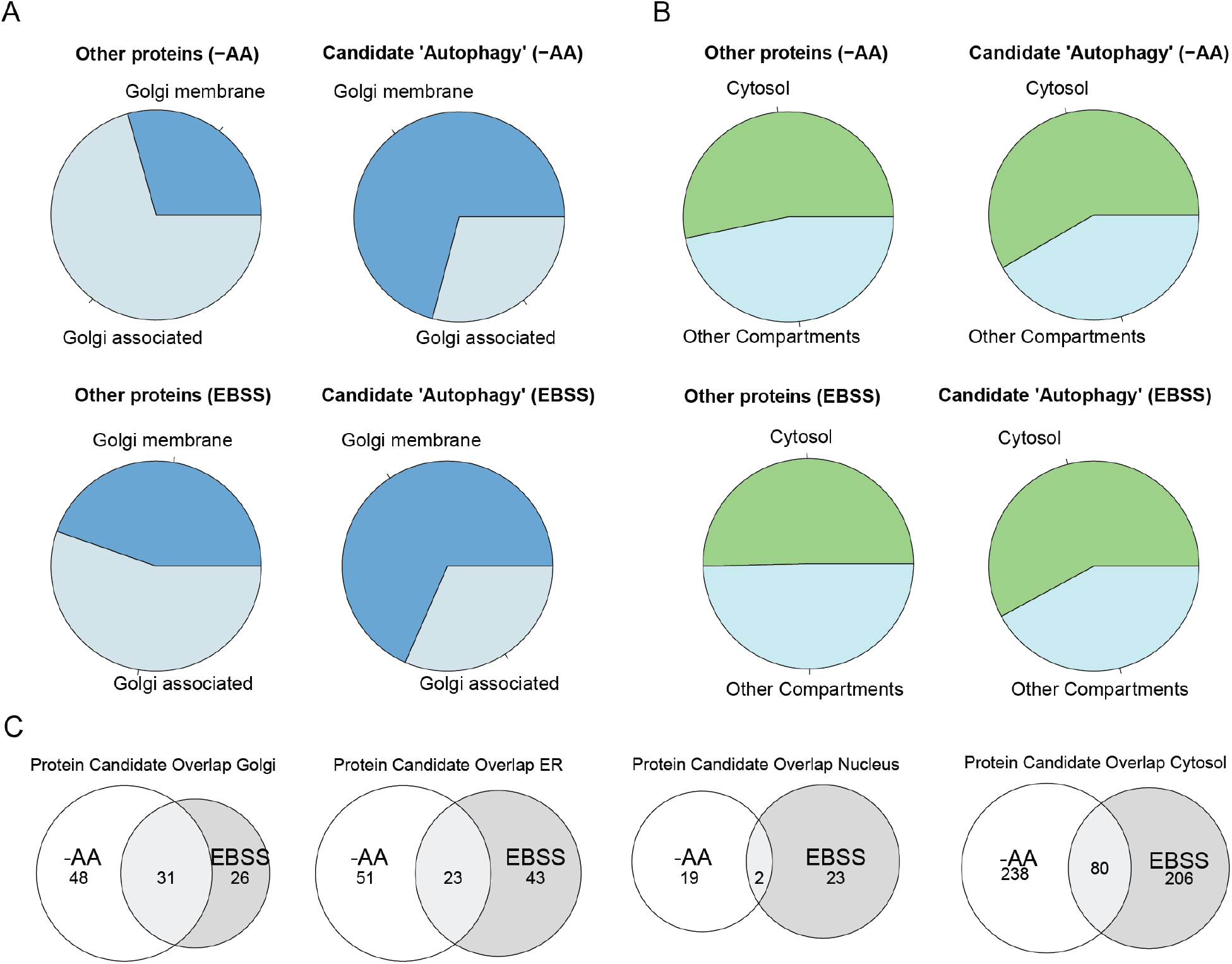
(**A**) Enrichment of Golgi-membrane and Golgi-associated proteins in the candidate ‘autophagy’ list and all other proteins for AA withdrawal and EBSS treatment. (**B**) Enrichment for the cytosolic proteins in the candidate ‘autophagy’ list and all other proteins for AA withdrawal and EBSS treatment. (**C**) Venn diagrams indicating the overlap of proteins identified in common within candidate ‘autophagy’ lists for AA withdrawal and EBSS treatment. Numbers within the diagram indicate the number of proteins present.

**fig. S7.**
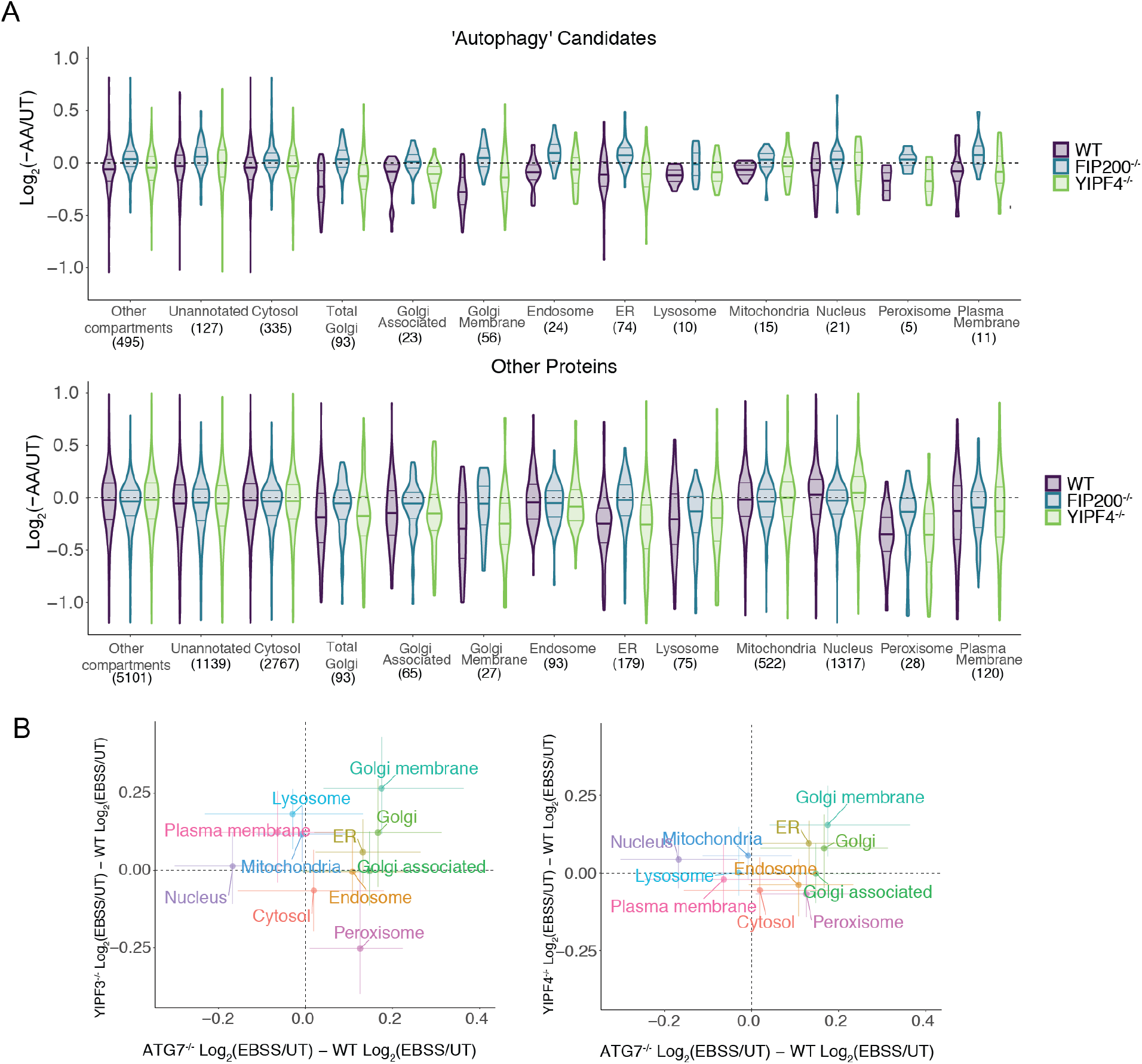
(**A**) Violin plots for Log_2_(-AA/UT) for control, FIP200^-/-^, or YIPF4^-/-^ HeLa cells displayed for various classes of proteins with the indicated sub-cellular localizations for either the ‘autophagy’ candidates or all other proteins from **fig. S5**. Median values are indicated by solid bold line. (**B**) Correlation plot for alterations in protein abundance for proteins in the indicated sub-cellular compartments in HeLa cells after 18 hours of EBSS for YIPF3^-/-^/WT or YIPF4^-/-^/WT cells (y-axis) versus FIP200^-/-^/WT cells (x-axis).

**fig. S8.**
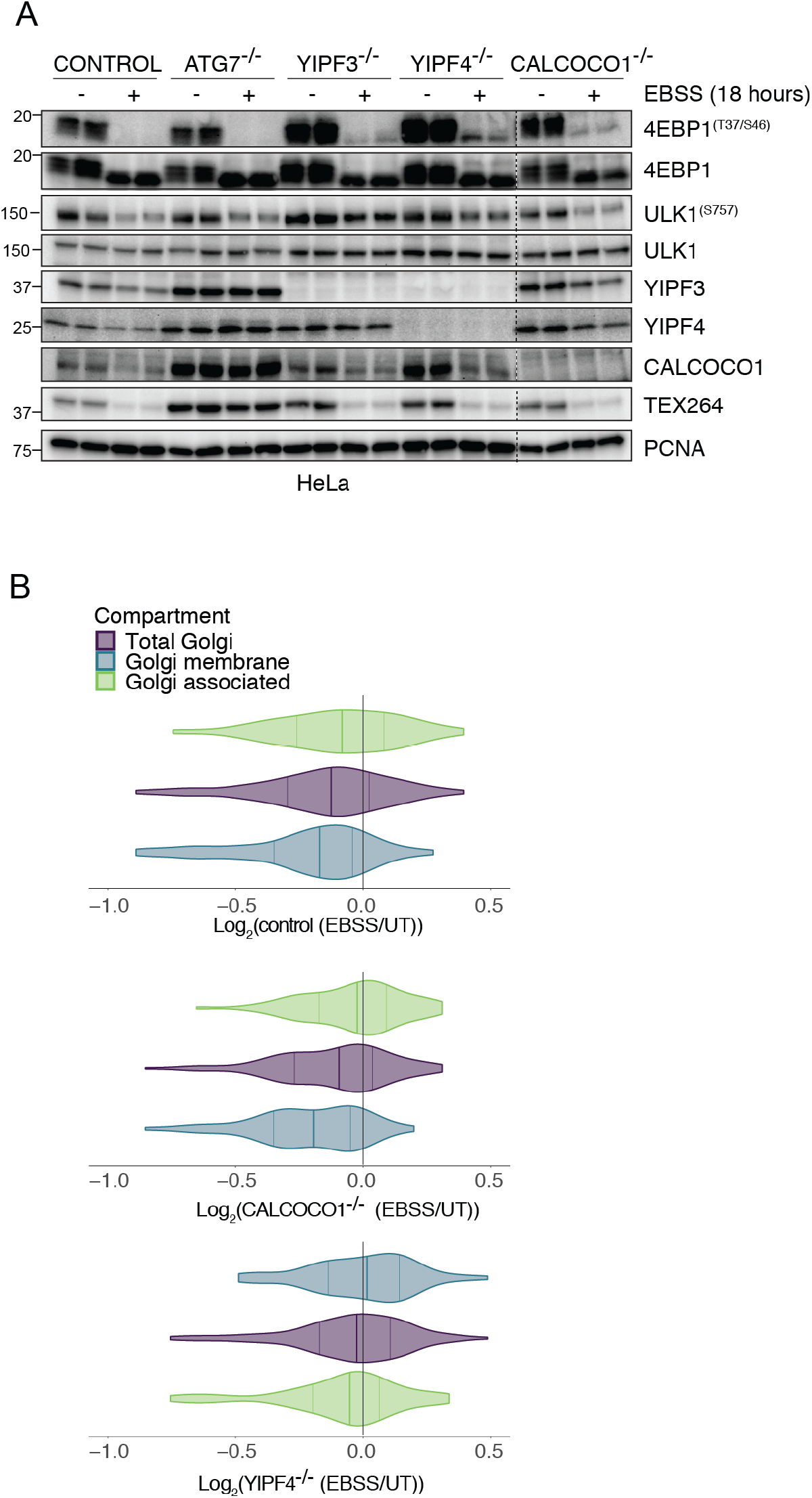
(**A**) Immunoblots of whole cell extracts from the indicated HeLa control and mutant cells in duplicate either left untreated or subjected to EBSS for 18 hours using the indicated antibodies. a-PCNA was used as a loading control. (**B**) Violin plot for Golgi-membrane protein Log_2_ FC with or without 18 hours of EBSS in control, YIPF4^-/-^ or CALCOCO1^-/-^ HeLa cells. Mean abundance is indicated by bold line.

**fig. S9.**
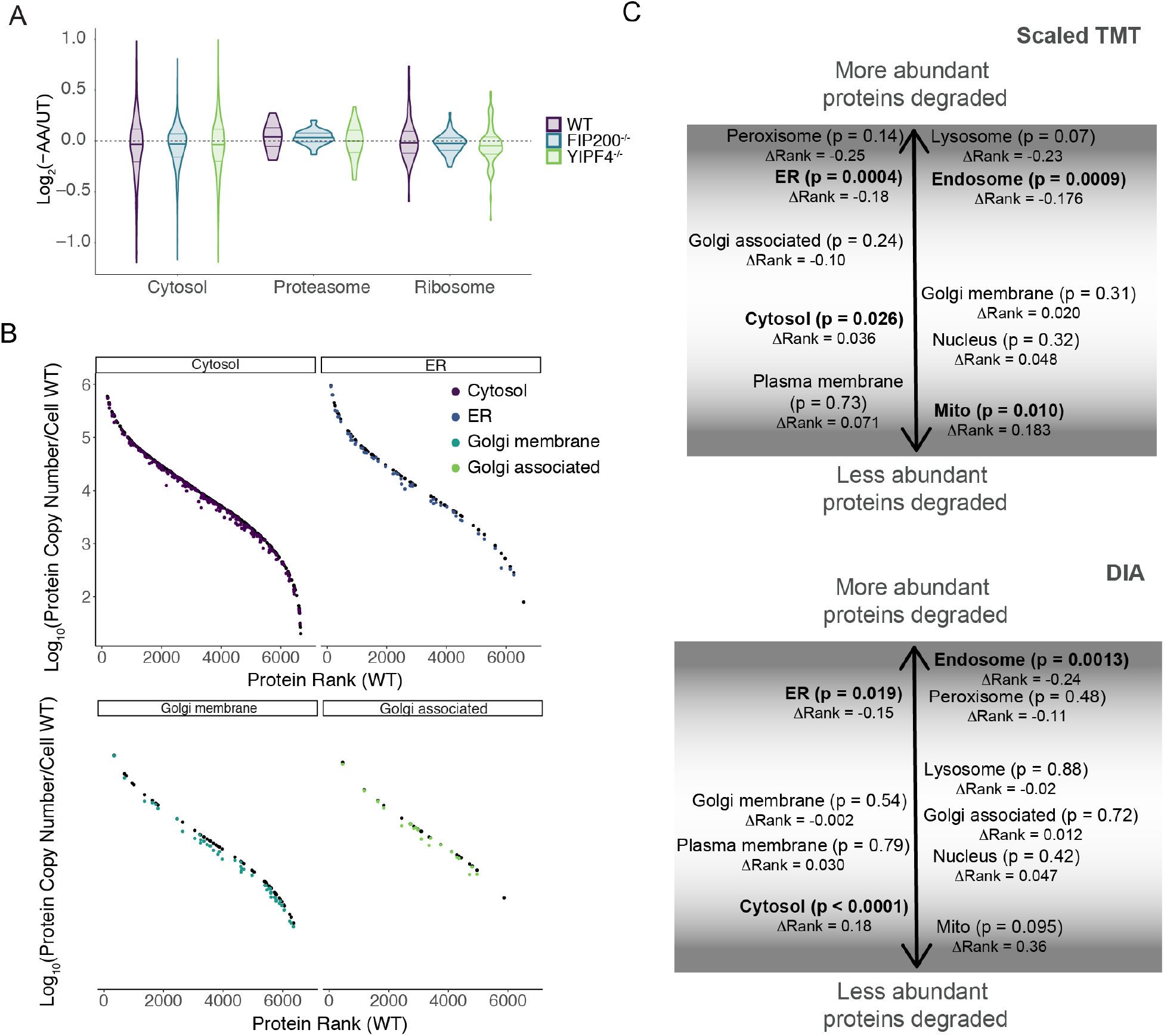
(**A**) Violin plots for Log_2_FC (-AA/UT) for control, FIP200^-/-^, or YIPF4^-/-^ HeLa cells displayed for 38 proteasome and 84 ribosomal proteins as well as proteins annotated as cytosolic. Median values are indicated by solid bold line. (**B**) Rank plot for cytoplasmic, ER and Golgi localized proteins. (**C**) Model for possible selectivity of macroautophagy at the organelle level. Abundance rank change (^ΔRank^) between proteins in the ‘autophagy’ candidate list – all other proteins for each organelle, scaled to number of total proteins in both scaled TMT and DIA experiments. For each compartment, p-values are listed and organelles with significant differences are in bold.

**fig. S10.**
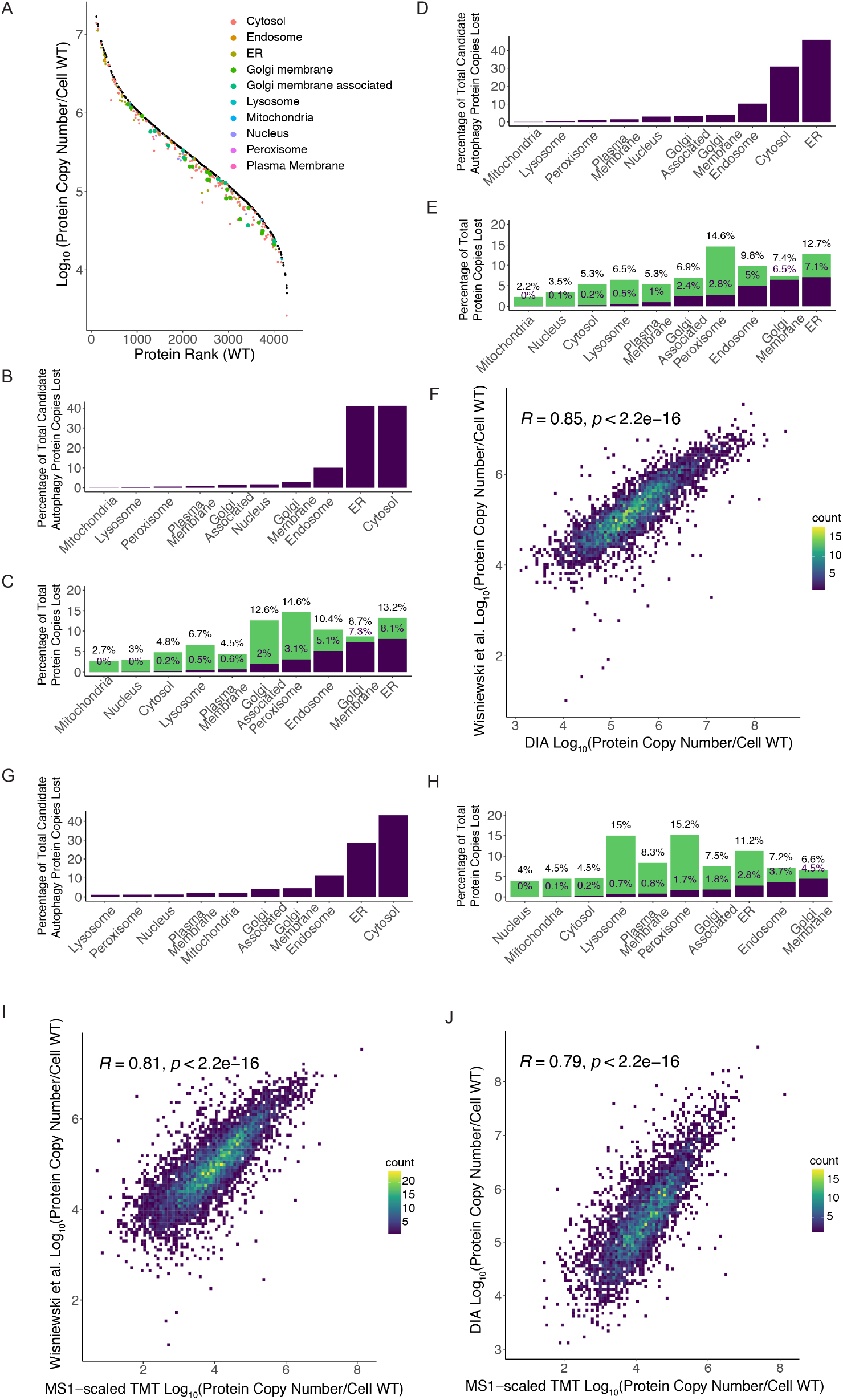
(**A**) DIA ranked plots. Protein copy number in the untreated condition for candidate ‘autophagy’ proteins in HEK293 cells (black) in rank order. The number of protein copies after loss by autophagy during amino acid starvation for each compartment as determined using protein abundance fold changes (AA withdrawal – untreated) by DIA. (**B**) Among the candidate autophagy proteins, percentage of total protein copy numbers lost via amino acid withdrawal (3.0161 x 10^7^ total). (**C**) Percentage of all protein copies lost from ‘autophagy’ candidate list (purple) or other mechanisms (green) by amino acid withdrawal for subcellular compartments based on DIA values with histone-based proteome ruler values. **(D-E**) Same as **B-C** respectively but based on DIA FC values mapped onto proteome ruler values from Wisniewski et al (*28*) (9.77 x 10^6^ total). (**F**) Correlation with DIA protein copy number estimates against Wisniewski et al (*28*) protein copy numbers. (**G-H**) Same as **B-C** respectively, based on TMT-scaled FC values mapped onto proteome ruler values from Wisniewski et al (*28*) (7.1573 x 10^6^ total). (**I**) Correlation plots for TMT-scaled MS1 protein signals against Wisniewski et al (*28*) copy number. (**J**) Correlation plots for TMT-scaled MS1 protein copy numbers and DIA protein copy numbers.

**fig. S11.**
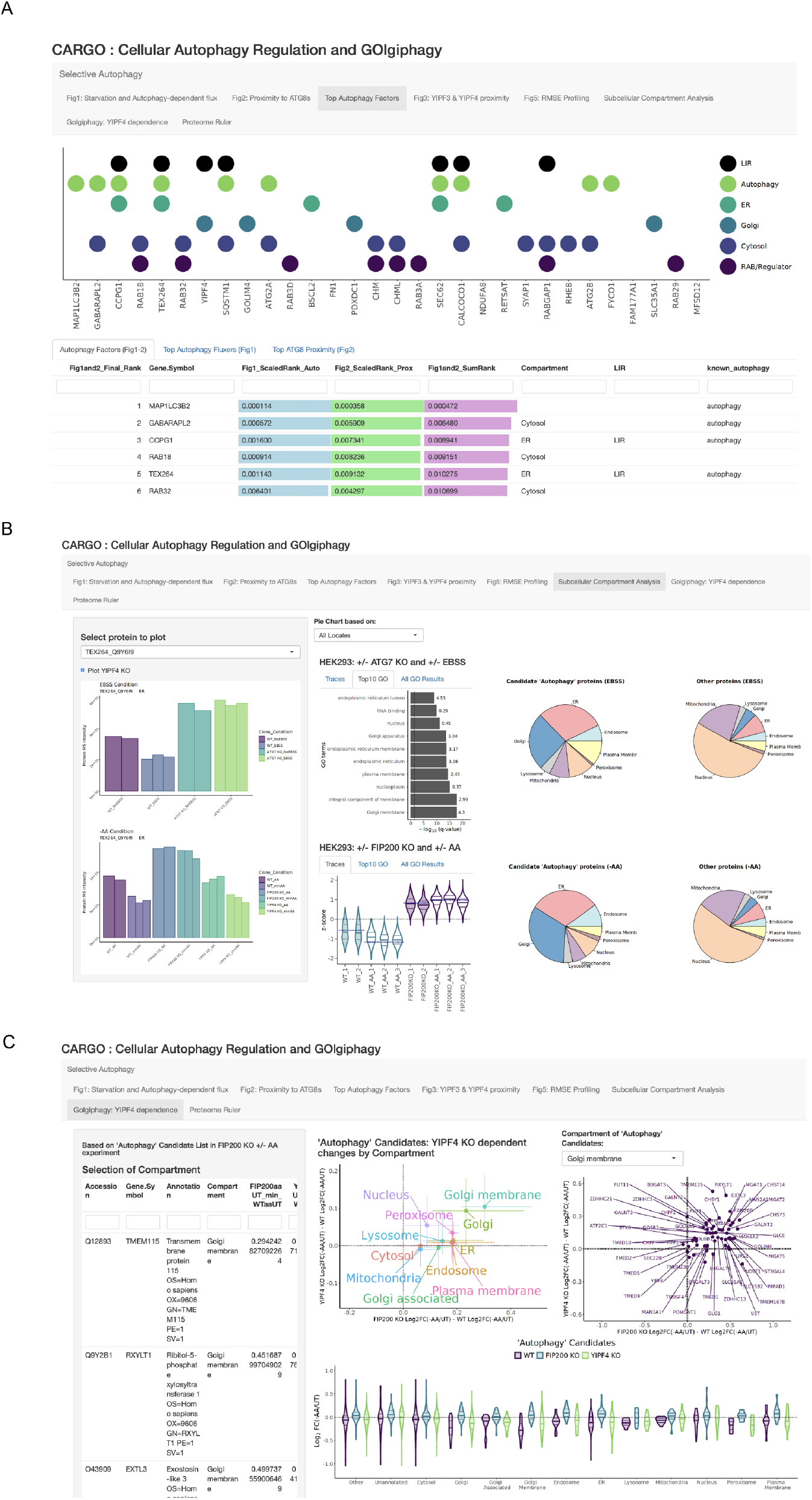
CARGO: an interactive website to interrogate Cellular Autophagy Regulation and Golgiphagy data from this work. The website can be found at: https://harperlab.connect.hms.harvard.edu/CARGO_Cellular_Autophagy_Regulation_GOlgiphagy/. (**A**) Example of visualization data combining Figure 1 and Figure 2 to create a priority list of putative autophagy factors. (**B**) Example of visualization data for Top Autophagy Fluxers and subcellular compartment analysis (Related to Figure 5). (**C**) Example of visualization tools for mapping Golgiphagy and autophagy clients.

